# Robust differential expression testing for single-cell CRISPR screens at low multiplicity of infection

**DOI:** 10.1101/2023.05.15.540875

**Authors:** Timothy Barry, Kaishu Mason, Kathryn Roeder, Eugene Katsevich

## Abstract

Single-cell CRISPR screens (perturb-seq) link genetic perturbations to phenotypic changes in individual cells. The most fundamental task in perturb-seq analysis is to test for association between a perturbation and a count outcome, such as gene expression. We conduct the first-ever comprehensive benchmarking study of association testing methods for low multiplicity-of-infection (MOI) perturb-seq data, finding that existing methods produce excess false positives. We conduct an extensive empirical investigation of the data, identifying three core analysis challenges: sparsity, confounding, and model misspecification. Finally, we develop an association testing method — SCEPTRE low-MOI — that resolves these analysis challenges and demonstrates improved calibration and power.

## Background

Pooled CRISPR screens with single-cell readout (e.g. Perturb-seq [1]) have emerged as a scalable, flexible, and powerful technique for connecting genetic perturbations to molecular phenotypes, with applications ranging from fundamental molecular biology to medical genetics and cancer research.[2] In such screens a library of genetic perturbations is transfected into a population of cells via CRISPR guide RNAs (gRNAs), followed by single-cell sequencing to identify the perturbations present and measure a rich molecular phenotype for each cell. The perturbations can target either genes [1] or non-coding regulatory elements,[3, 4, 5] either repressing [1] or activating [6] these targets; the molecular readouts can include gene expression,[1] protein expression,[7, 8, 9] or epigenetic phenotypes like chromatin accessibility.[10] Typically, perturbations are introduced at low multiplicity of infection (MOI), with one perturbation per cell. In cases where perturbations are expected to have weak effects (like regulatory-element-targeting screens), perturbations also can be introduced at high-MOI (with many perturbations per cell) to increase scalability.[3, 4, 5, 11, 12]

The most fundamental statistical task involved in the analysis of single-cell CRISPR screen data is to test for association between a perturbation and a univariate, count-based molecular phenotype, such as the expression of a gene or protein. In our previous work on high-MOI single-cell CRISPR screen analysis, we discovered that existing methods for association testing are prone to an excess of false positive hits.[13] In that work we proposed SCEPTRE, a well-calibrated method for association testing on high-MOI data. Since low-MOI screens currently outnumber high-MOI screens, the low-MOI association testing problem is even more pressing. A variety of methods has been deployed for association testing in low-MOI.[1, 14, 15, 16, 9, 8, 17, 10] However, there is no consensus as to which of these methods represents the “state of the art;” these methods have not undergone rigorous statistical validation and comparison;[18] and in fact there is no commonly accepted framework for quantifying the statistical validity of single-cell CRISPR screen association testing methods. Resolving these fundamental issues is essential to ensuring the reliability of biological conclusions made on the basis of single-cell CRISPR screen experiments.

We aimed to address the aforementioned challenges through three contributions. First, we developed a simple framework for evaluating the calibration of association testing methods for single-cell CRISPR screens. We then leveraged this framework to conduct the first-ever [18] comprehensive benchmarking study of association methods on low-MOI data, applying six leading methods to analyze six diverse datasets. We found that all existing methods exhibit varying degrees of miscalibration, indicating that results obtained using these methods may be contaminated by excess false positive discoveries. Second, to shed light on why existing methods might demonstrate miscalibration, we conducted an in-depth empirical investigation of the data, uncovering three core analysis challenges: confounding, model misspecification, and data sparsity. No existing method addresses all of these analysis challenges, explaining their lack of calibration. Finally, we developed SCEPTRE (low-MOI), a substantial extension of the original SCEPTRE [13] tailored to the analysis of low-MOI single-cell CRISPR screens. SCEPTRE (low-MOI) is based on the novel and statistically principled technique of permuting negative binomial score statistics. (We often will refer to the low-MOI version of SCEPTRE simply as “SCEPTRE” for the sake of brevity). SCEPTRE addresses all three core analysis challenges both in theory and in practice, demonstrating markedly improved calibration and power relative to existing methods across datasets. SCEPTRE is available at katsevich-lab.github.io/sceptre/.

## Results

### A survey of leading analysis methods

Association testing on low-MOI single-cell CRISPR screens is a variation on the classical single-cell differential expression testing problem (Figure 1a). To test for association between a given targeting CRISPR perturbation and gene, one first divides the cells into two groups: those that received the targeting perturbation, and those that received a non-targeting (NT) perturbation. (All other cells typically are ignored.) One then tests for differential expression of the given gene across these two groups of cells, yielding a fold change estimate and *p*-value. One repeats this procedure for a (typically) large, preselected set of perturbation-gene pairs. Finally, one computes the discovery set by subjecting the tested pairs to a multiplicity correction procedure (e.g., Benjamini-Hochberg).

**Figure 1.**
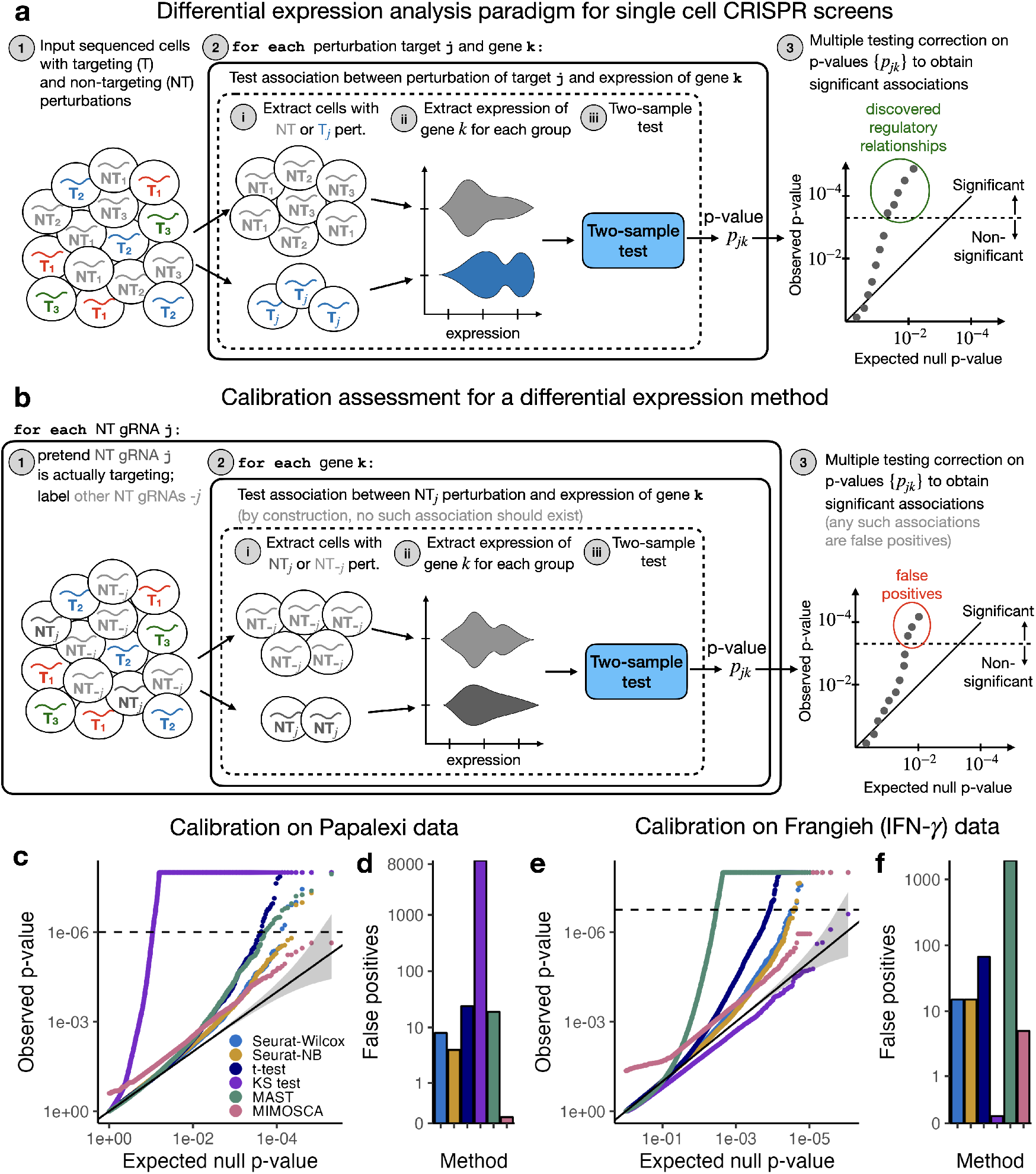
Comprehensive benchmarking study of single-cell CRISPR screen association testing methods on low-MOI data. **a**, The standard paradigm for association testing on low-MOI single-cell CRISPR screen data. To test for association between a given targeting perturbation and gene, one tests for differential expression of the gene across two groups of cells: those containing the given targeting perturbation, and those containing a non-targeting (NT) perturbation. One typically repeats this procedure for a large, preselected set of targeting-perturbation gene pairs, obtaining a discovery set by subjecting the resulting *p*-values to a multiple comparison correction procedure (e.g., Benjamini-Hochberg). **b**, The calibration check paradigm. One constructs “null” or “negative control” perturbation-gene pairs by coupling each individual NT gRNA to the entire set of genes. One then assesses the calibration of a method by deploying the method to analyze these null pairs. Any *p*-values that survive the multiple testing correction procedure correspond to false positive discoveries. **c-d**, Results of the calibration check benchmarking analysis on the Papalexi gene expression data. **c**, QQ plot of the null *p*-values (colored by method) plotted on a negative log transformed scale. Gray region, 95% confidence band. **d**, Number of false discoveries that each method makes on the null pairs after a Bonferroni correction at level 0.1. **e-f**, Similar to panels **c-d**, but for the Frangieh IFN-*γ* data.

We use the term “control group” to refer to the cells against which the cells that received the targeting perturbation are compared. As indicated above, the control group typically is the set of cells that received an NT perturbation (i.e., the “NT cells”). Certain single-cell CRISPR screen methods, however, take as their control group the set of cells that did *not* receive the targeting perturbation (i.e., the “complement set”). In low-MOI screens the NT cells generally constitute a more natural control group than the complement set, as we seek to compare the effect of the targeting perturbation to that of a “null” perturbation rather than to the average of the effects of all other perturbations introduced in the pooled screen. In high-MOI screens, however, the complement set is the only choice, because few (if any) cells receive only NT perturbations.

We surveyed recent analyses of single-cell CRISPR screen data and identified five methods commonly in use: the default Seurat [19] `FindMarkers()` function based on the Wilcoxon test (Seurat-Wilcox); MIMOSCA;[1] a *t*-test on the library-size-normalized expressions;[10] MAST;[20] and a Kolmogorov-Smirnov (KS) test on the library-size-normalized expressions.[21] We also considered applying `FindMarkers()` with negative binomial (NB) regression rather than the Wilcoxon test (Seurat-NB). These methods vary along several dimensions (Table 1; Existing methods details), including their testing paradigm (two-sample test versus regression-based test), how they normalize the data, whether they make parametric assumptions, and whether they use the NT cells or the complement set as their control group. Most of these methods are popular single-cell differential expression procedures that have been adapted to single-cell CRISPR screen setting.

**Table 1:** A summary of low-MOI single-cell CRISPR screen DE methods currently in use. The applications of each method to single-cell CRISPR screens are cited below the method name. The methods vary along several key axes, including the use (or lack thereof) of parametric assumptions, the construction of the null distribution, the variables adjusted for, and the control group. NT, non-targeting.

| Method | Paradigm | Parametric assumption | Null distribution | Normalization/ Adjustments | Control group |
| --- | --- | --- | --- | --- | --- |
| Seurat-Wilcox [7, 9, 22] | Two-sample test | No | Asymptotic | Library size | NT cells |
| MIMOSCA[1] [1, 23, 24, 25, 8, 26] | Regression-based | No | Permutation | Library size, other covariates | Complement set |
| $t$-test [10] | Two-sample test | Yes | Asymptotic | Library size | NT cells |
| MAST[20] [15] | Regression-based | Yes | Asymptotic | Library size, expressed genes | NT cells |
| KS test [21, 16] | Two-sample test | No | Asymptotic | Library size, batch | NT cells |
| Seurat-NB (single-cell DE) | Two-sample test | Yes | Asymptotic | Library size | NT cells |

### Comprehensive benchmarking study of leading analysis methods

We sought to assess whether these methods are correctly calibrated (i.e., whether they yield uniformly distributed *p*-values under the null hypothesis of no association between the perturbation and gene). Methods that are not correctly calibrated can produce discovery sets that are contaminated by excess false positives or false negatives. Unfortunately, there does not exist a standard protocol for assessing the calibration of single-cell CRISPR screen association methods. The closest existing analysis [15] proceeds by applying methods to analyze gene-perturbations pairs for perturbations with known targets. Any pair where the gene is not the known target of the perturbation is considered null. As acknowledged by the original authors, this approach underestimates precision because downstream effects of perturbations are not taken into account.

To help fill this methodological gap, we designed a simple procedure to ascertain the calibration of a single-cell CRISPR screen association method (Figure 1b). We constructed a set of “null” or “negative control” perturbation-gene pairs by pairing each NT gRNA to each gene. We then deployed a given method to analyze these null pairs. (For methods that use the NT cells as their control group — the majority of methods — this check consists of comparing cells containing a given NT gRNA to cells containing *all other* NT gRNAs.) The output of this check is a set of *N*_gene_ *· N*_NT_ null *p*-values, where *N*_gene_ is the number of genes and *N*_NT_ is the number of NT gRNAs. Since the null perturbation-gene pairs are devoid of signal, a well-calibrated association method should output uniformly distributed *p*-values on these pairs. Deviations from uniformity — and thus miscalibration of the method — can be detected by inspecting a QQ plot of the *p*-values. Quantitatively, the number of null pairs passing a Bonferroni correction measures the extent of the miscalibration; well-calibrated methods should have roughly zero such pairs.

We note that there are two uses of the proposed calibration check procedure. The first use is for the goal of benchmarking existing analysis methods to identify which, if any, are suitable for broad application (the primary goal of this section). The second use is for the goal of testing whether a *given* method is well-calibrated on a *given* dataset. These two goals are distinct; a method may not be broadly well-calibrated but may perform adequately on a given dataset. In the context of the second goal, we recommend applying a modified calibration check where the set of negative control perturbation-gene pairs is matched to the set of pairs under consideration based on several criteria (described later).

We employed the above framework to systematically benchmark the performance of the existing methods, implementing each as faithfully as possible in a publicly available R package `lowmoi` (github.com/katsevich-lab/lowmoi). We applied the calibration check procedure using six single-cell CRISPR screen datasets, five real and one simulated (Additional file 1: Tables S1-S2). The five real datasets came from three papers: Frangieh 2021 [8] (three datasets), Papalexi 2021 [9] (one dataset), and Schraivogel 2020 [15] (one dataset). The data were diverse, varying along the axes of CRISPR modality (CRISPRko or CRISPRi), technology platform (perturb-CITE seq, ECCITE-seq, or targeted perturb-seq), cell type (TIL, K562, or THP1), and genomic element targeted (enhancers or gene TSS). Notably, the Papalexi data were multimodal, containing both gene and protein expression measurements. For simplicity we analyzed the gene and protein modalities separately throughout.

Surprisingly, the results of our analyses (Figure 1c-f; Additional file 1: Figures S1, S2, S3) revealed substantial miscalibration for many dataset-method pairs. On the Papalexi data, for example, the KS test produced inflated *p*-values, yielding over 9,000 false Bonferroni discoveries. MAST was similarly inflated on the Frangieh IFN-*γ* data, falsely rejecting nearly 2,000 null perturbation-gene pairs. MIMOSCA, meanwhile, exhibited noticeably non-uniform behavior on both datasets, outputting *p*-values strictly less than 0.26 across all pairs. Overall, the two best methods appeared to be Seurat-Wilcox and Seurat-NB, although these two methods still demonstrated clear signs of miscalibration. We noted that the calibration quality of a given method could vary significantly across datasets; this is explained by the fact that different datasets posed different analysis challenges. Nevertheless, we concluded that none of the methods was adequately calibrated across all datasets tested, suggesting that existing methods may not be suitable for broad application to single-cell CRISPR screen data.

### Systematic identification of core analysis challenges

We conducted an extensive empirical investigation of the data to search for possible sources of miscalibration, uncovering three core analysis challenges: sparsity, confounding, and model misspecification. No method that we examined addressed more than one of these analysis challenges (Table S3), explaining their collective lack of calibration. This section is abbreviated to preserve space; interested readers can consult Details of the investigation into the core analysis challenges, which contains a more detailed description of this investigation.

Single-cell CRISPR screen data typically are sparse, both in terms of gene expression and perturbation presence. Many genes have nonzero expression in only a small fraction of cells. On the other hand, due to the pooling of a large number of perturbations in a single experiment, the perturbation presence data are also sparse: most perturbations are present in only a small fraction of cells. The latter sparsity distinguishes single-cell CRISPR screens from other single-cell applications and is particularly pronounced in low-MOI. To summarize both sources of sparsity in a single number, we defined the “effective sample size” for a given perturbation-gene pair as the number of cells containing both the perturbation and nonzero gene expression.

We found that effective sample size had a substantial effect on the calibration of many methods under consideration (Additional file 1: Figures S4-S5), especially those based on asymptotic approximations, such as Seurat-Wilcox. Asymptotic approximations tend to break down when the effective sample size is too low. For example, we compared the exact null distribution of the Wilcoxon test statistic (obtained via permutations) to the asymptotic Gaussian distribution used by Seurat-Wilcox; the latter is a computationally tractable approximation to the former in large samples. The Gaussian distribution provided a reasonable approximation to the exact null distribution for some pairs (Figure 2a, left) but not others (2a, right). Furthermore, as the effective sample size decreased and the Gaussian approximation degraded in accuracy, the *p*-value obtained via the Gaussian approximation likewise degraded in accuracy (Figure 2b). Finally, stratifying the Seurat-Wilcox null *p*-values by effective sample size on the Frangieh IFN-*γ* data revealed that pairs with small effective sample sizes yielded more inflated *p*-values than pairs with large effective sample sizes (Figure 2c).

**Figure 2.**
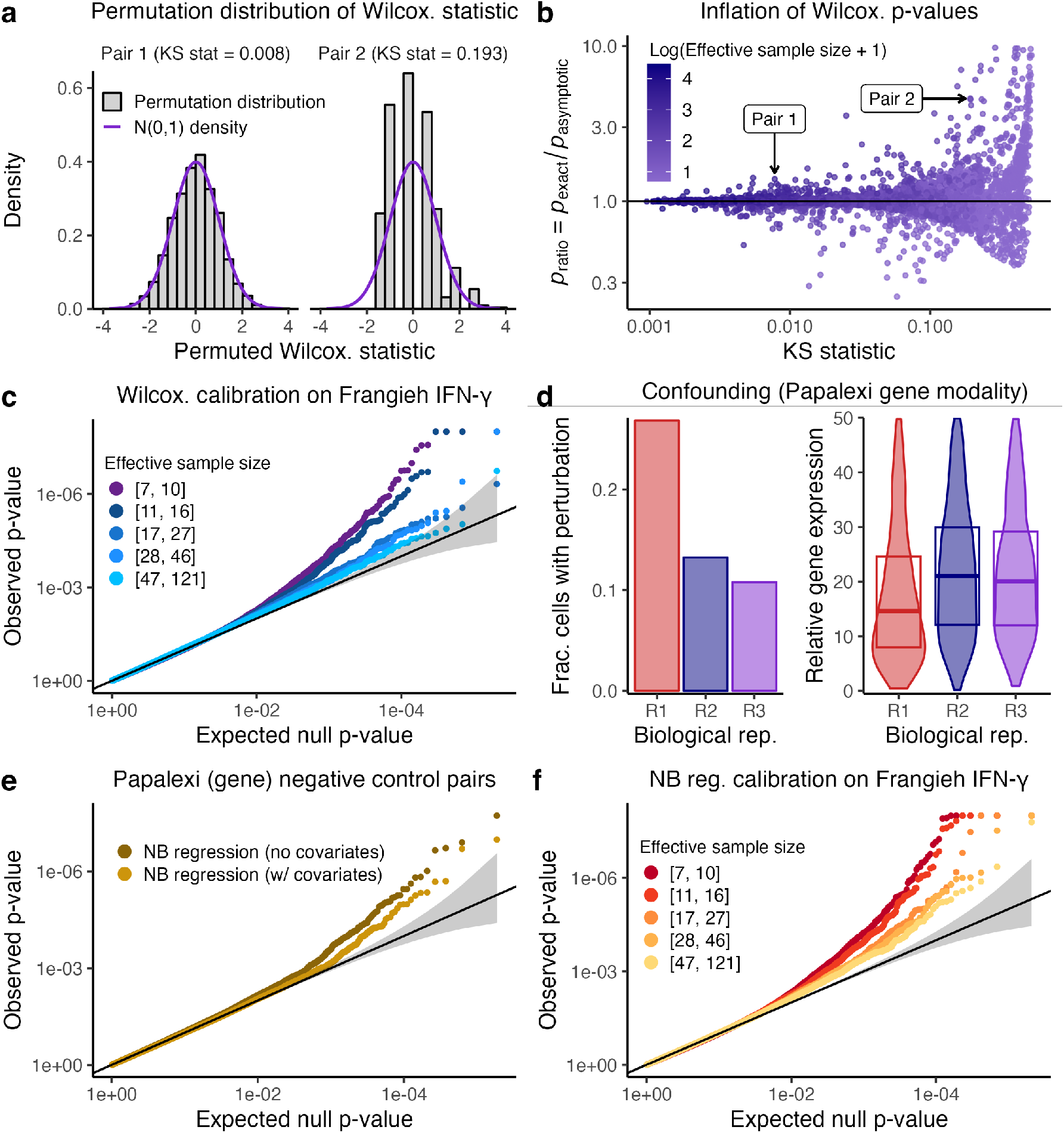
Sparsity, confounding, and model misspecification are core analysis challenges in single-cell CRISPR screen analysis. **a**, The exact null distribution of the Wilcoxon test statistic (obtained via permutations; gray) on two pairs from the Frangieh IFN-*γ* data. The Wilcoxon test (and thus Seurat-Wilcox) approximates the exact null distribution using a standard Gaussian density (purple). For pair 1 (left), the Gaussian approximation to the exact null distribution is good (goodness of fit KS statistic = 0.008), while for pair 2 (right) the approximation is inadequate (goodness of fit KS statistic = 0.193). **b**, A plot of *p*_ratio_ (defined as the ratio of the exact Wilcoxon *p*-value, *p*_exact_, to the asymptotic Wilcoxon *p*-value, *p*_asymptotic_) vs. goodness of fit of the Gaussian distribution to the exact null distribution (as quantified by the KS statistic). Each point represents a gene-gRNA pair; pairs 1 and 2 (from panel **a**) are annotated. As the KS statistic increases (indicating worse fit of the Gaussian distribution to the exact Wilcoxon null distribution), *p*_ratio_ deviates more from one, indicating miscalibration. Points are colored according to the effective sample size of the corresponding pair. **c**, Stratification of the Seurat-Wilcox *p*-values on the Frangieh IFN-*γ* negative control data by effective sample size. **d**, An example of confounding on the Papalexi data. Left (resp. right), the fraction of cells that received a given NT gRNA (resp., the relative expression of a given gene) across biological replicates “R1,” “R2,” and “R3.” **e**, Application of NB regression with and without covariates to the Papalexi data. **f**, Stratification of the NB regression *p*-values on the Papalexi (gene expression) negative control data by effective sample size.

Second, technical factors, such as biological replicate, batch, and library size, impact not only a cell’s expression level, but also its probability of receiving a perturbation, thereby creating a confounding effect that can lead to spurious associations [13]. All existing methods adjust for library size, but few adjust for other technical factors (Table 1). We studied how the variable of biological replicate confounded the Papalexi (gene modality) data (Figure 2d). The Papalexi data were sequenced across three separate biological replicates (which we labeled “R1,” “R2,” and “R3”). We visually examined the relationship between biological replicate and a given NT gRNA (“NTg4”) and gene (*FTH1*). We plotted the fraction of cells in each biological replicate that harbored “NTg4” (Figure 2d, left); additionally, we plotted the (library-size-normalized) expression level of *FTH1* across biological replicate, superimposing boxplots indicating the 25th, 50th, and 75th percentiles of the distribution (Figure 2d, right). We observed clear visual evidence that biological replicate impacted *both* NTg4 presence or absence *and FTH1* expression level, creating a confounding effect. For example, cells in biological replicate R1 were much more likely than cells in biological replicates R2 or R3 to contain NTg4; on the other hand, cells in biological replicate R1 exhibited a much lower expression level of *FTH1* than cells in biological replicates R2 or R3. If one naively tested for association between NTg4 and *FTH1* while ignoring batch, one would incorrectly conclude that NTg4 *decreased* the expression of *FTH1*. In fact, NTg4 exerted no effect on the expression *FTH1*, as NTg4 was a negative control gRNA.

To assess the utility of adjusting for technical factors beyond library size, we applied negative binomial (NB) regression — both with and without biological replicate included as a covariate — to the Papalexi negative control data (Figure 2e). The variant of NB regression with biological replicate, though not perfectly calibrated, outperformed its counterpart without biological replicate. Methods not adjusting for biological replicate on the Papalexi data (such as Seurat-Wilcox) exhibited *worse* calibration for large effective sample sizes (Additional file 1: Figures S4-S5), where there is more power to detect the spurious confounding-driven associations.

Third, methods that rely upon parametric models for the gene expression distribution, such as NB regression and MAST, can yield miscalibrated *p*-values when those models are misspecified.[27] To assess this effect, we monitored *p*-value calibration of the NB regression method on the Frangieh IFN-*γ* data while gradually increasing the effective sample size (Figure 2f). We found that the calibration quality improved until a point before plateauing; even for large effective sample sizes, noticeable miscalibration remained. (The non-parametric Seurat-Wilcox method, by contrast, attained good calibration for large effective sample sizes on this dataset.) This pattern was consistent with poor fit of the NB regression model, potentially due to inadequate estimation of the NB size parameter.

### SCEPTRE (low-MOI) addresses the analysis challenges

We next developed SCEPTRE (low-MOI), a method for robust single-cell CRISPR screen association testing on low-MOI data (Figure 3a). For a given targeting perturbation-gene pair, SCEPTRE first regresses the vector of gene expressions onto the vector of perturbation indicators and matrix of technical factors via an NB GLM. (A given entry of the perturbation indicator vector is set to “1” if the corresponding cell contains a targeting perturbation and “0” if it contains a non-targeting perturbation.) SCEPTRE then computes the *z*-score *z*_*obs*_ for a test of the null hypothesis that the the coefficient corresponding to the perturbation indicator in the fitted GLM is zero. Next, SCEPTRE permutes the perturbation indicator vector *B* times (while holding fixed the gene expression vector and technical factor matrix) and recomputes a *z*-score for each of the permuted indicator vectors, yielding *B* “null” *z*-scores. Finally, SCEPTRE fits a smooth (skew-normal) density to the histogram of null *z*-scores and computes a *p*-value by evaluating the tail probability of the fitted density based on the original test statistic *z*_obs_.

**Figure 3.**
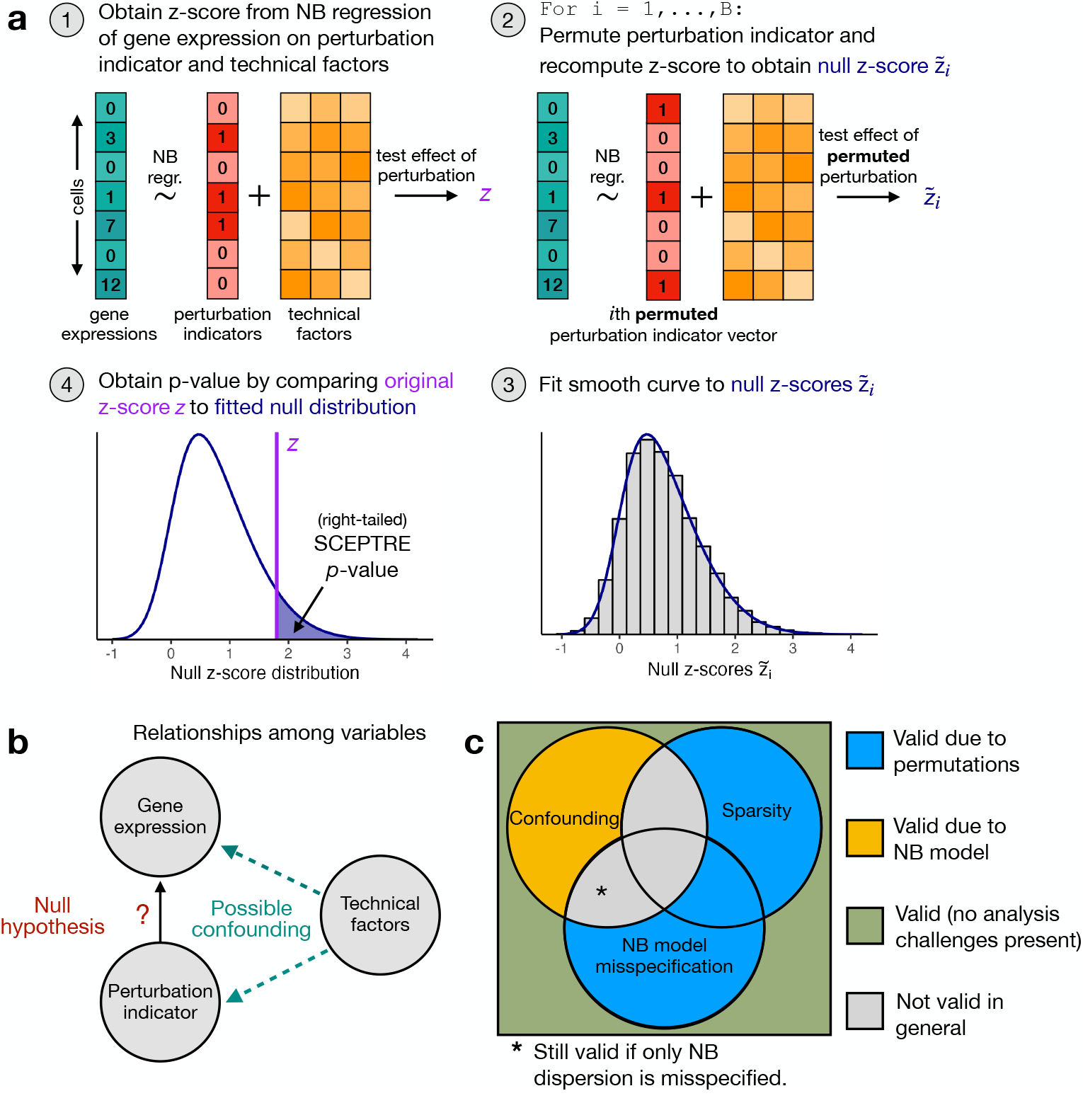
SCEPTRE addresses the core analysis challenges of sparsity, confounding, and model misspecification in theory. **a**, The SCEPTRE algorithm. First, the gene expressions are regressed onto the perturbation indicators and technical factors, and the *z*-score *z*_obs_ corresponding to the perturbation indicator is computed. Second, the perturbation indicators are permuted (while the gene expressions and technical factors are held fixed) and the *z*-score is recomputed, yielding *B* “null” *z*-values. Third, a smooth density is fit to the histogram of the null *z*-values. Fourth, a *p*-value is computed by evaluating the tail probability of the fitted density at *z*_obs_. **b**, A diagram representing the relationship between the variables in the analysis. The technical factors often (but not always) exert a confounding effect on the perturbation indicator and gene expression. **c**, A diagram illustrating the robustness properties of SCEPTRE. The circles represent analysis challenges. A perturbation-gene pair can be affected any subset of the analysis challenges. The color in each region of the diagram indicates whether SCEPTRE is valid on pairs affected by that subset of analysis challenges (blue, yellow, or green = valid; gray = not valid in general). For regions in which SCEPTRE is valid, the color of the region indicates *why* SCEPTRE is valid (yellow = NB model, blue = permutations). The validity of SCEPTRE is overdetermined on pairs unaffected by *any* analysis challenge (green region) due to the combination of the NB model and permutations.

SCEPTRE possesses several appealing theoretical and computational properties. Theoretically, SCEPTRE is robust to the calibration threats of sparsity, confounding, and model misspecification. A key observation is that the technical factors (e.g., biological replicate) may or may not exert a confounding effect on the perturbation indicator and gene expression (Figure 3b). If confounding is absent for a given perturbation-gene pair, then SCEPTRE is valid even when the NB model is misspecified or problem is highly sparse. On the other hand, if confounding is present, then SCEPTRE retains validity if the NB model is correctly specified and the problem is not too sparse (Figure 3c). (In fact, in the latter case, empirical results indicate that the NB model need only be specified correctly up to the dispersion parameter, sidestepping the difficult problem of NB dispersion parameter estimation.[28, 29]) In this sense SCEPTRE is the only method that addresses all three core analysis challenges (Additional file 1: Table S3). We empirically demonstrated the above key robustness property of SCEPTRE in simulation experiments (Additional file 1: Figure S7 and S11).

SCEPTRE also is performant, capable of analyzing hundreds of perturbation-gene pairs per second. We attained this efficiency by implementing several computational accelerations. First, we elected to use a score test (as opposed to a more standard Wald or likelihood ratio test) to compute the NB *z*-scores; the score test enabled us to fit a single NB GLM per perturbation-gene pair and share this fitted GLM across all permuted perturbation indicator vectors. Second, we derived a new algorithm for computing GLM score tests, which is hundreds of times faster than the classical algorithm when the perturbation indicator vector is sparse, as is often the case in single-cell CRISPR screen analysis (Additional file 1: section “Comparing the spectral decomposition algorithm to the QR decomposition algorithm for computing GLM score tests”). Finally, we developed a novel strategy — “inductive without replacement sampling” — for recycling compute across permutation tests in which each test contains the same number of control units (Additional file 1: section “Inductive without replacement sampling”).

A natural question is whether SCEPTRE is better (in some sense) than the simpler method that entails “regressing out” the technical factors via an NB GLM, extracting the residuals from the fitted GLM, and then performing a permutation test on the residuals (taking, for example, the difference in means across treatment and control groups as the test statistic). We answered this question in the affirmative, finding that SCEPTRE was considerably more powerful than the alternative, residual-based method on both real and simulated data (Additional file 1: section “Comparing the score statistic to the difference-in-residual-means statistic”). Finally, we note that SCEPTRE (low-MOI) is inspired by, but distinct in several ways from SCEPTRE (high-MOI).[13] We clarify similarities and differences between these two methods in Comparison of SCEPTRE (low-MOI) and SCEPTRE (high-MOI).

### Application of SCEPTRE to negative and positive control data

We included SCEPTRE in the calibration benchmarking analysis presented before. An inspection of the QQ plots revealed that SCEPTRE markedly improved on the calibration of the two best existing methods, namely Seurat-Wilcox and Seurat-NB (Figure 4a-b). For example, on the Frangieh IFN-*γ* data, SCEPTRE made one Bonferroni rejection and yielded *p*-values that lay mostly within the gray 95% confidence band. The Seurat methods, by contrast, made fifteen false rejections each and produced *p*-values that fell considerably outside the confidence band. Next, we tabulated the number of Bonferroni-significant false positives for each dataset-method pair (Figure 4c; smaller values are better). SCEPTRE generally made the fewest number of false discoveries among all methods. On average over datasets, SCEPTRE made only 0.7 false discoveries, a roughly tenfold improvement over the Seurat methods.

**Figure 4.**
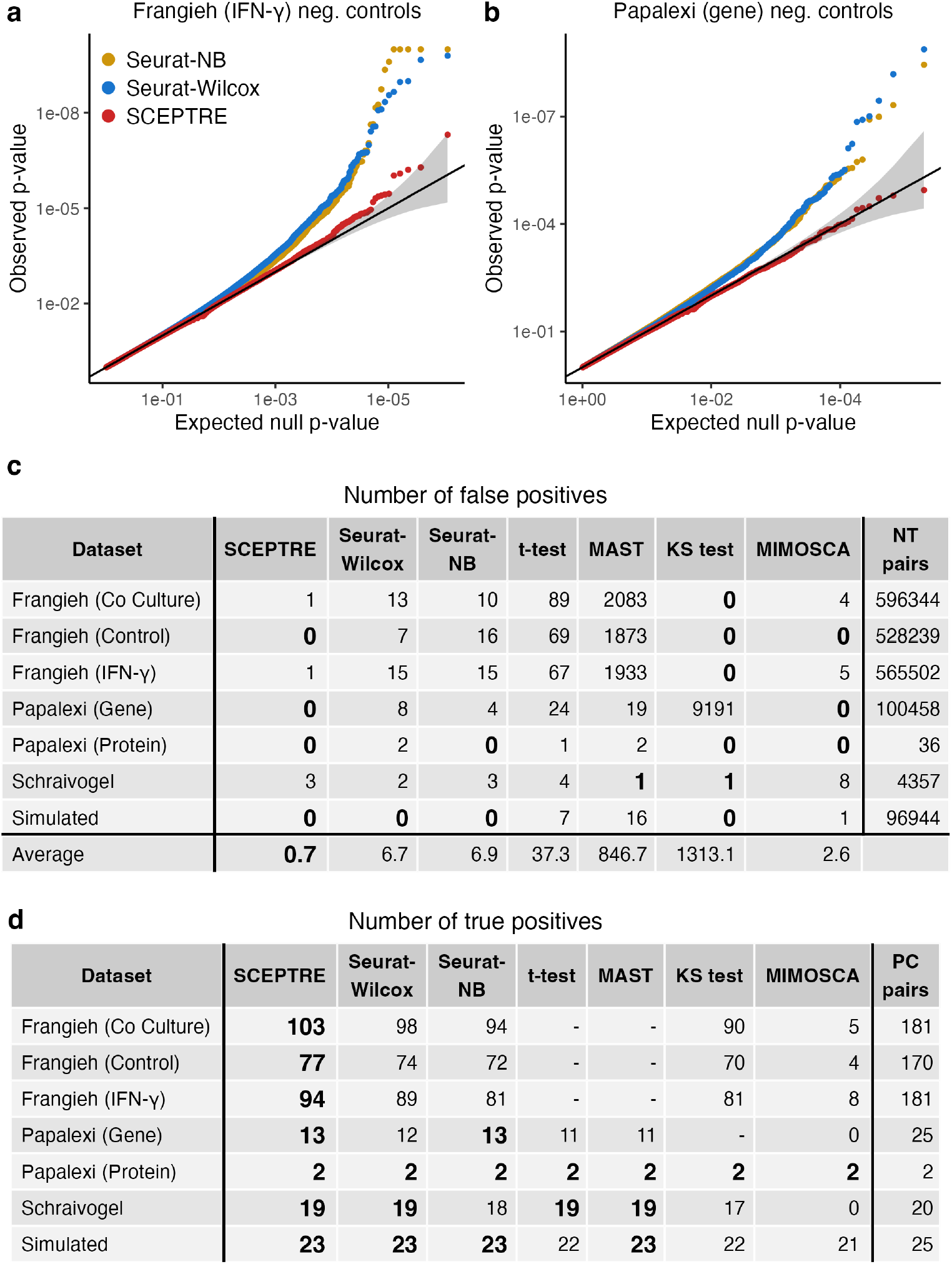
SCEPTRE demonstrates improved calibration and power relative to existing methods across datasets. **a** (resp. **b**), QQ plot of the *p*-values outputted by Seurat-NB, Seurat-Wilcox, and SCEPTRE on the Frangieh IFN-*γ* (resp., Papalexi gene expression) negative control data. Gray band, 95% confidence region. **c**, Number of false discoveries (at Bonferroni correction level 0.1) on the negative control data for each method-dataset pair. **d**, Number of true discoveries (significant at level *α* = 10^*−*5^) on the positive control data for each each method-dataset pair.

Next, we assessed the power of the methods by applying them to positive control data. We constructed positive control pairs for each dataset by coupling perturbations targeting TSSs to the genes (or proteins) regulated by these TSSs. We examined the number of “highly significant” discoveries — operationally defined as rejections made at level *α* = 10^*−*5^ — made by each method on each dataset (Figure 4d; larger values are better). Methods that exhibited extreme miscalibration on a given dataset (defined as *>* 50 Bonferroni rejections on the negative control pairs of that dataset) were excluded from the positive control analysis, as assessing the power of such methods is challenging. We found that SCEPTRE matched or outperformed the other methods with respect to power on every dataset (while at the same time achieving better calibration on negative control data).

### Pairwise quality control and experimental design

Quality control (QC) — the removal of low-quality cells — is a key step in the analysis of single-cell data. In the context of single-cell CRISPR screens, it is useful not only only to remove low-quality cells but also low-quality perturbation-gene pairs. We term this latter type of QC “pairwise QC.” As discussed previously, effective sample size — the number of cells containing both the perturbation and nonzero gene expression — affects the calibration of several methods considered. It also affects power, as small effective sample sizes yield low power and therefore needlessly increase the multiplicity burden. We found that SCEPTRE rarely rejected positive control pairs with an effective sample size below seven (Additional file 1: Figure S8); moreover, SCEPTRE maintained calibration for negative control pairs with an effective sample size of seven and above (Additional file 1: Figures S4-S5). For this reason our pairwise QC strategy consisted of filtering for pairs with an effective sample size of seven or greater. We applied this pairwise QC throughout.

Additionally, we reasoned that our results on the power and calibration of SCEPTRE might inform questions related to the experimental design of single-cell CRISPR screens, such as the number of cells per perturbation required to ensure adequate power and calibration of SCEPTRE on a given dataset. We derived a simple mathematical expression for the minimum number of cells that must contain each perturbation so as to ensure that a specified fraction of pairs passes pairwise QC, where the pairwise QC threshold was selected on the basis of SCEPTRE’s calibration and power at different effective sample sizes. We concluded that, on a dataset of standard sparsity (e.g., the Frangieh IFN-*γ* dataset or the Papalexi gene modality dataset), each perturbation should be contained within at least 50-65 cells to ensure that 95% of pairs pass pairwise QC. (See Additional file 1: section “Experimental design considerations” for more details.)

### Application of SCEPTRE for discovery analyses

The standard workflow involved in applying SCEPTRE to analyze a new single-cell CRISPR screen dataset consists of three main steps. First, the user prepares the data to pass to SCEPTRE and defines the “discovery set,” which is the set of perturbation-gene pairs that the user seeks to test for association. (A reasonable default choice is the set of all possible pairs.) Second, the user runs the “calibration check” to verify that SCEPTRE is adequately calibrated on the dataset under analysis. The calibration check involves applying SCEPTRE to analyze a set of automatically-constructed negative control pairs. These negative control pairs are “matched” to the discovery pairs in several respects. For example, the negative control pairs and discovery pairs are subjected to the exact same pairwise QC, and the number of negative control pairs is set equal to the number of discovery pairs. If the calibration check fails, the user can take steps to improve calibration, such as adding covariates or varying the QC thresholds. After verifying adequate calibration, the user runs the “discovery analysis,” which entails applying SCEPTRE to analyze the pairs contained in the discovery set (Figure 5a).

**Figure 5.**
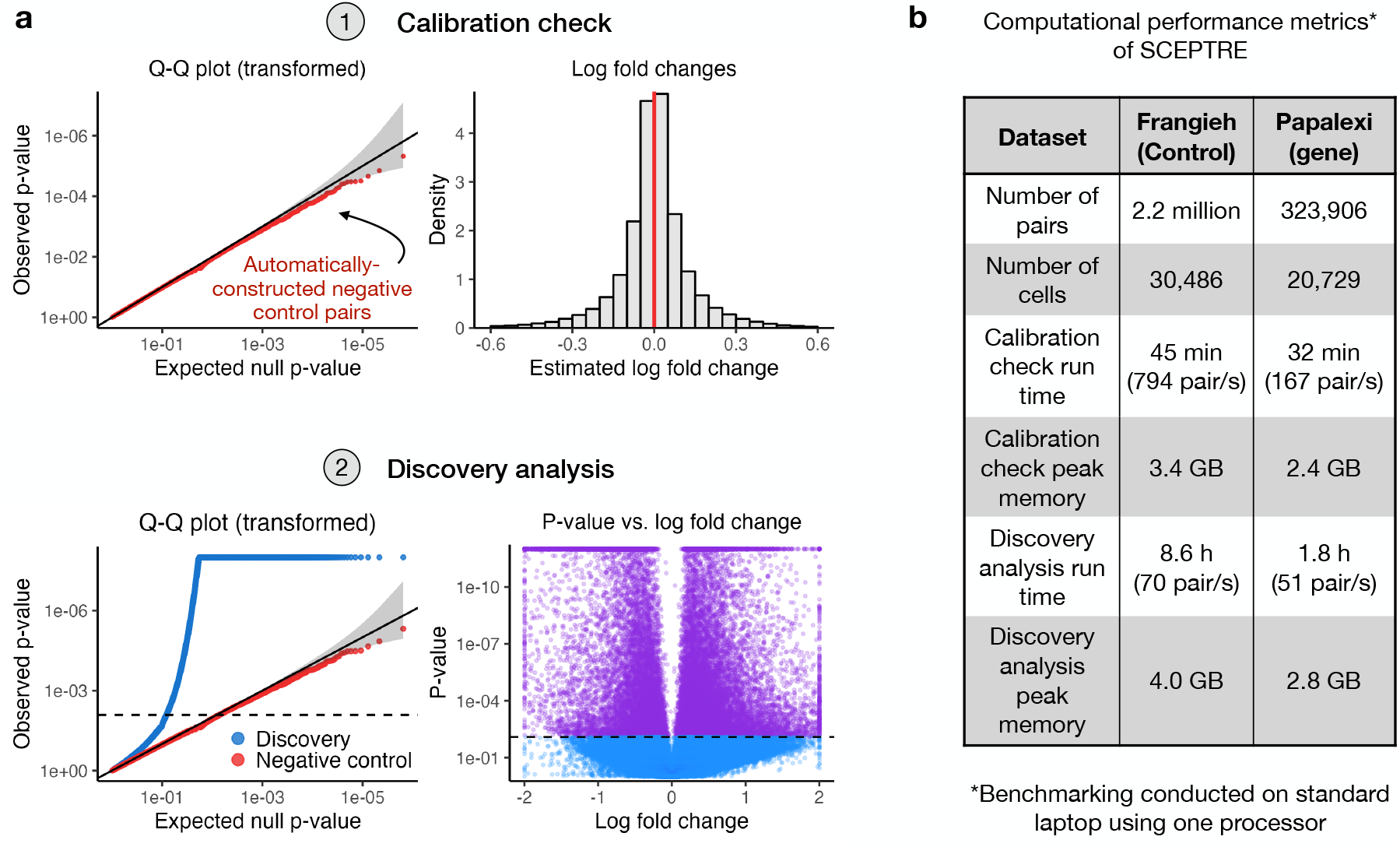
Applying SCEPTRE to make biological discoveries. **a**, The standard workflow involved in applying SCEPTRE to a new dataset, using the Papalexi gene expression data as a running example. First, SCEPTRE is applied to analyze a set of automatically-constructed negative control pairs (the “calibration check”). The resulting negative control *p*-values are plotted on a QQ plot to ensure uniformity (upper left), and the negative control log-fold changes are plotted on a histogram to ensure symmetry about zero (upper right). Second, SCEPTRE is applied to analyze the discovery pairs (the “discovery analysis”). The discovery *p*-values are superimposed over the negative control *p*-values to ensure that signal is present in the discovery set (lower left), and a volcano plot is created (lower right). **b**, Computational performance metrics of SCEPTRE on the Frangieh (control) and Papalexi (gene expression) data. A complete *trans* analysis was conducted on both datasets. Several metrics are reported, including calibration check run time, calibration check peak memory usage, discovery analysis run time, and discovery analysis peak memory usage.

To illustrate the above workflow, we applied SCEPTRE to carry out a complete *trans* analysis of the Papalexi (gene expression) and Frangieh (control) datasets. Many of the genes targeted for knockout in these datasets were transcription factors (TFs); thus, our main biological objective was to map the TFs to their target genes. We carried out a calibration check and discovery analysis on both datasets (Figure 5b). These fairly large analyses completed within a matter of hours on a single laptop processor and required a few gigabytes of memory.

To validate the results of the discovery analysis, we identified the downstream target genes of the transcription factors STAT1 and IRF1 using cell-type-relevant ChIP-seq data.[30] We examined the degree of concordance between the discovery set produced by each method and the ChIP-seq-identified targets (Additional file 1: Figure S9). To this end, for each method, we constructed a two-by-two contingency table of genes contained within the discovery set of the method and genes whose TSS overlapped with a ChIP-seq peak. We computed a p-value (via a Fisher exact test) on this contingency table, quantifying the extent to which the method’s discovery set was enriched for ChIP-seq signal.

We made several observations. First, the discovery set of the KS test did not demonstrate enrichment for ChIP-seq signal (enrichment *p >* 0.5), likely because the KS test produced a large number of false discoveries. Second, and somewhat surprisingly, MIMOSCA’s discovery set exhibited the greatest degree of enrichment for ChIP-seq signal (among all methods) on STAT1 and the lowest degree of enrichment (among all methods, excluding the KS test) on IRF1. However, MIMOSCA made many fewer discoveries than any other method on the discovery data (in particular, 956 and 875 fewer discoveries than the next closest method for IRF1 and STAT1), a trend consistent with MIMOSCA’s results on the the positive control data^1^ (Figure 4). Among the methods that made a large number of discoveries on both the positive control data and the discovery data (i.e., all methods except for MIMOSCA), SCEPTRE ranked second out of six for ChIP-seq signal enrichment on both IRF1 and STAT1, with the top method being different across the two transcription factors. Thus, SCEPTRE appeared to exhibit consistently good performance on the discovery data (and more broadly across all of our analysis; Figure 4), increasing our confidence in the results.

## Discussion

Single-cell CRISPR screens have emerged as a powerful method for linking genetic perturbations to rich phenotypic profiles in individual cells. Although poised to impact a variety of research areas, single-cell CRISPR screens will play an especially important role in dissecting the regulatory logic of the noncoding genome. The bulk of genetic risk for diseases lies in noncoding regions, implicating dysregulation of gene expression.[31, 32, 33] A major challenge in genetics, therefore, is to map noncoding disease variants to the genes that they target, target genes to the molecular programs that they regulate, and — ultimately — molecular programs to disease.[34] Single-cell screens have enabled breakthrough progress on these tasks. For example, two recent studies leveraged high-MOI single-cell screens to perturb blood disease [11] and cancer [35] GWAS variants (in some cases at single nucleotide resolution) and link these variants to target genes in disease-relevant cell types. (Both studies used SCEPTRE (high-MOI) to analyze their data.) Another recent study leveraged low-MOI single-cell screens to knock down genes regulated by heart disease GWAS variants and map these genes to downstream molecular programs.[34] Given the promise that single-cell screens have demonstrated in understanding noncoding variation, a wave of single-cell screens aiming to link noncoding variants to genes and genes to molecular programs likely will emerge over the coming decade.

It is therefore crucial that reliable methods for single-cell CRISPR screen data analysis be made available. The broad objective of this work was to put single-cell CRISPR screen analysis onto a more solid statistical foundation. To this end we devised a simple framework for assessing the calibration and power of competing methods; applied this framework to conduct the first-ever comprehensive benchmarking study of existing methods; identified core statistical challenges that the data pose; and developed a method, SCEPTRE, that combines careful modeling with a resampling framework to produce a well-calibrated, powerful, fast, and memory-efficient test of association. Taken together, these contributions help bring statistical rigor to single-cell CRISPR screen data analysis. Furthermore, given the appealing theoretical properties and empirical performance of the proposed method, we anticipate that the method could be extended (with some modification) to applications beyond single-cell CRISPR screens, such as single-cell eQTL analysis and single-cell case-control differential expression analysis.[36]

We identified sparsity, confounding, and model misspecification as key challenges in single-cell CRISPR screen analysis. However, the data pose additional challenges that SCEPTRE does not currently address. First, some NT gRNAs may have off-targeting effects. In such cases testing for association by comparing cells that contain a targeting perturbation to those that contain an NT perturbation could result in a loss of error control. At least one prior work has attempted to resolve this problem.[37] Second, some targeting gRNAs are ineffective, i.e., they fail to perturb their target. Including such defective gRNAs in the analysis can result in a loss of power. Several methods, including MIMOSCA,[1] MUSIC,[38] and Mixscape,[9] have been developed to address this issue. Third, it is challenging to distinguish between direct and indirect effects in the sense that perturbations can be associated with their direct targets or with targets further downstream. Disentangling direct from indirect effects likely admits a statistical solution, but to our knowledge, this problem remains unaddressed. Finally, genes often are co-expressed in “gene modules.” An exciting opportunity is to pool information across genes within the same module to increase the power of perturbation-to-gene association tests; the recent method GSFA does this in a Bayesian framework.[39]

## Conclusions

Single-cell CRISPR screens are a promising technology for functional genomics discovery. However, the analysis of single-cell CRISPR screen data presents several statistical and computational challenges, demanding the development of new analytic methods. The SCEPTRE toolkit, which now supports both low- and high-MOI CRISPR screens, provides practitioners with a unified solution for statistically reliable and computationally efficient single-cell CRISPR screen differential expression analysis.

## Methods

### Dataset details

We downloaded, processed, and harmonized five single-cell CRISPR screen datasets (Additional file 1: Table S1), inheriting several data-related analysis decisions made by the original authors. First, we used the gRNA-to-cell assignments that the original authors used, thereby circumventing the need to assign gRNAs to cells using gRNA UMI and/or read count matrices. Papalexi and Schraivogel employed a simple strategy for this purpose: Papalexi identified the gRNA with the greatest UMI count in a given cell and assigned that gRNA to the cell, while Schraivogel assigned gRNAs by thresholding gRNA UMI counts. Frangieh, meanwhile, assigned gRNAs to cells via a more complex approach involving a separate dial-out PCR procedure. We found the gRNA-to-cell assignments adequate and thus used them without modification. Next, we inherited the cell-wise QC that the original authors implemented. For example, Papalexi removed likely duplets (as determined by the `Seurat` function `MULTIseqDemux`[40, 41]) as well as cells with excessive mitochondrial content and low gene expression.

We generated a synthetic, negative control single-cell CRISPR screen dataset to use for bench-marking the calibration of the competing methods. The synthetic dataset contained 5,000 genes, 25 gRNAs, and 10,000 cells. We generated the matrix of gene expressions by sampling counts from a negative binomial distribution, allowing each gene to have its own mean and size parameter. (We drew gene-wise means and sizes i.i.d. from a Gamma(0.5, 2) distribution and a Unif(1, 25) distribution, respectively.) We randomly inserted gRNAs into cells such that the expected number of cells per gRNA was equal across gRNAs. The dataset was entirely devoid of signal and confounding: no gRNA affected the expression of any gene, and no technical factors impacted the gRNA assignments or gene expressions. We also generated a synthetic positive control dataset to assess the power of the competing methods under known ground truth. The synthetic positive control dataset contained 125 genes, 25 positive control gRNAs, 100 negative control gRNAs, and 15,000 cells. The mean expression of each gene across “treatment” and “control” groups was drawn (separately) from a Gamma(0.5, 2) distribution. The gene-wise sizes were drawn from a Unif(1, 25) distribution. Like the negative control dataset, the positive control dataset was devoid of confounding.

We applied our own minimal gene-wise, gRNA-wise, and cell-wise QC uniformly to the datasets. We filtered for genes expressed in at least 0.005 of cells, gRNAs expressed in at least 10 cells, and cells with exactly one gRNA, respectively. Table S2 summarizes the statistical attributes (e.g., number of genes, number of cells, etc.) of each dataset. Finally, we obtained the set of cell-specific covariates (or technical factors) for each dataset, which we list below. Frangieh co-culture, control, and IFN-*γ* datasets: number of gene UMIs, number of genes expressed; Papalexi (gene modality): number of gene UMIs, number of genes expressed, biological replicate, and percent of gene transcripts that mapped to mitochondrial genes; Papalexi (protein modality): number of protein UMIs, biological replicate, and percent of gene transcripts that mapped to mitochondrial genes; Schraivogel: number of gene UMIs, number of genes expressed, sequencing lane.

### Existing methods details

We benchmarked the performance of six methods: Seurat-Wilcox, Seurat-NB, *t*-test, MAST, KS test, and MIMOSCA. The first five of these methods are generic single-cell differential expression methods that have been adapted to single-cell CRISPR screens (either by us or other single-cell researchers), while MIMOSCA is specific to single-cell screens. To facilitate benchmarking of the methods, we implemented all in an R package `lowmoi` (github.com/Katsevich-Lab/lowmoi). We implemented Seurat-Wilcox and Seurat-NB via a call to the Seurat `FindMarkers()` function. In the case of Seurat-Wilcox, we called `NormalizeData()` before `FindMarkers()` to normalize the gene expressions by dividing the gene expressions by library size. Next, we implemented the *t*-test via a call to `t.test()` in R. Following Liscovitch et al.,[10] we normalized the gene expression vector for a given gene-perturbation pair by dividing by the library size, subtracting the mean, and dividing by the standard deviation. We used the implementation of MAST that Schraivogel et al. used to analyze their single-cell screen data.[15] To this end we copied and pasted relevant portions of the Schraivogel et al. Github codebase (github.com/argschwind/TAPseq_manuscript) into `lowmoi`. Similarly, we used the implementation of the KS test that Replogle et al. used to analyze their single-cell screen data,[16] again copying and pasting relevant portions of the corresponding codebase into `lowmoi` (github.com/thomasmaxwellnorman/Perturbseq_GI). Finally, we implemented MIMOSCA by copying and pasting relevant sections of the MIMOSCA package (github.com/klarman-cell-observatory/Perturb-CITE-seq) into `lowmoi`. Replogle et al.’s implementation of the KS test and MIMOSCA both were written in Python. Thus, we used the `reticulate` package to access these methods from within R. To ensure consistency of the API across methods, we implemented the methods in such a way that each took the same inputs and returned the same output. Finally, to ensure correctness, we tested for agreement between the output of our implementations and those of the original methods (when possible).

Some methods have an internal QC step in which gene-perturbation pairs that are unpromising or low-quality (as determined by the method itself) are removed. For example, Seurat DE by default filters out gene-perturbation pairs for which the log-fold change of the expression of the gene (across the treatment and control cells) falls below a certain threshold. We disabled such method-specific pairwise QC, allowing us to apply competing methods to the exact same set of gene-perturbation pairs on each dataset, facilitating head-to-head comparisons across methods.

We applied several variants of NB regression to the data. First, as described above, we applied Seurat-NB the negative control and positive control pairs of all datasets. Furthermore, as part of our investigation into the analysis challenges (section Systematic identification of core analysis challenges), we applied NB regression as implemented by the `MASS`[42] package to the Papalexi (gene expression) and Frangieh IFN-*γ* negative control data. (These results are depicted in Figure 2e-f). We used the `MASS` implementation of NB regression in exploring the analysis challenges, as `MASS` is slightly more flexible than Seurat, in particular enabling the straightforward inclusion of covariates. Within the context of `MASS` NB regression, we tested for association between a perturbation and the expression of a gene via a GLM score test, as implemented by the `statmod`[43] package. We elected to use a score test (as opposed to a more standard Wald or likelihood ratio test) test to make our implementation of NB regression more comparable to SCEPTRE, as SCEPTRE uses a permutation test built upon an NB regression score test statistic.

### Details of the calibration check procedure

We describe the calibration check procedure in greater detail. Suppose there are *d* distinct NT gRNAs; index these gRNAs from 1 to *d*. Let 𝒞_1_ denote the set of cells containing NT gRNA 1, 𝒞_2_ the set of cells containing NT gRNA 2, etc. Let 𝒞 = 𝒞_1_ ∪ 𝒞_2_ ∪ · · · ∪ 𝒞_*d*_ denote the set of cells containing *any* NT gRNA (i.e., the “NT cells”). Next, let *𝒞 \ 𝒞*_*i*_ denote the set of cells containing *any* NT gRNA *excluding* NT gRNA *i*. Additionally, let *𝒯* denote the set of cells containing *any targeting* gRNA. (Observe that *𝒯 ∪ 𝒞* is the set of all cells.) Finally, let *𝒯 ∪ 𝒞 \ 𝒞*_*i*_ denote the set of all cells excluding the cells that contain NT gRNA *i*. Let there be *p* distinct genes.

Suppose we seek to check the calibration of a given method. The way in which we deploy the method to analyze a given negative control pair depends on whether the method uses the NT cells or the complement set as its control group (Table 1). Consider the negative control pair formed by coupling NT gRNA *i* to gene *j*. If the method uses the NT cells as its control group (e.g., Seurat-Wilcox, Seurat-NB, SCEPTRE, etc.), then we apply the method to test for differential expression of gene *j* across the groups of cells 𝒞_*i*_ and *𝒞\𝒞*_*i*_. By contrast, if the method uses the complement set as its control group (e.g., MIMOSCA), then we apply the method to test for differential expression of gene *j* across the groups of cells *𝒯 ∪ 𝒞 \ 𝒞*_*i*_ and 𝒞_*i*_. The *effective sample size* of the given negative control pair is the number of cells in the set 𝒞_*i*_ for which the expression of gene *j* is nonzero. In carrying out our benchmarking analysis (Figure 1c-f, Figure 4), we restricted our attention to the subset of the *d · p* possible negative control pairs whose effective sample size was greater than or equal to seven.

For testing calibration on a given input dataset, the SCEPTRE software automatically constructs a set of negative control pairs that is matched to the pairs in the “discovery set” — i.e., the set of targeting perturbation-gene pairs that the user seeks to test for association — in several respects. First, the negative control pairs and discovery pairs are subjected to the same pairwise QC. Second, the number of negative control pairs is set equal to the number of discovery pairs (assuming the number of possible negative control pairs matches or exceeds the number of discovery pairs). Third, if the user elects to “group” together gRNAs that target the same site (as opposed to running an analysis in which singleton gRNAs are tested for significance), then the negative control pairs likewise are constructed by “grouping” together individual NT gRNAs. Overall, the negative control pairs are designed to mirror the discovery pairs, the difference being that the negative control pairs are devoid of biological signal.

### Details of the investigation into the core analysis challenges

We describe in greater detail our empirical investigations into the core analysis challenges of sparsity, confounding, and model misspecification (as described in Section Systematic identification of core analysis challenges).

#### Sparsity

To explore the impact of sparsity on calibration, we deployed the two-sample Wilcoxon test to a randomly-selected subset of 5,400 negative control gene-gRNA pairs from the Frangieh IFN-*γ* data. (The pairs were selected such that each had an effective sample size of one or greater.) Following Seurat-Wilcox, we deployed the Wilcoxon test as follows: first, we normalized the gene expressions by dividing the raw counts by the cell-specific library sizes; then, we applied the Wilcoxon test (as implemented by the `wilcox.test` function from the `stats` package in R) to the normalized data, comparing the treatment cells to the control cells. Finally, we computed the Wilcoxon *p*-value in two ways. First, we calculated the asymptotic *p*-value *p*_asymptotic_ by comparing the Wilcoxon test statistic to the standard Gaussian distribution. This approach implicitly assumes that the number of cells with nonzero expression (across both groups) is large enough for the null distribution of the Wilcoxon test statistic to be approximately Gaussian. Next, we calculated the exact *p*-value *p*_exact_ by (i) computing the Wilcoxon statistic on the original data; (ii) permuting the gRNA indicator vector *B* = 200, 000 times (while holding fixed the vector of normalized gene expressions), resulting in *B* permuted datasets; (iii) computing the Wilcoxon test statistic on each of these *B* permuted datasets, yielding a permutation (or “null”) distribution of Wilcoxon statistics; and then (iv) calculating the *p*-value *p*_exact_ by comparing the original Wilcoxon statistic to the null Wilcoxon statistics[44]. The latter approach, though computationally expensive (due to the slowness of computing the Wilcoxon statistic), yields a much more accurate *p*-value than the asymptotic approach for lowly expressed genes. Seurat-Wilcox returns the asymptotic *p*-value *p*_asymptotic_ instead of the exact *p*-value *p*_exact_ in virtually all cases.^2^

To study the impact of making the above approximation, we plotted the asymptotic null distribution of the Wilcoxon statistic (i.e., the standard Gaussian distribution) superimposed on top of the exact null distribution of the Wilcoxon statistic (i.e., the permutation distribution) for two pairs from the Frangieh IFN-*γ* negative control data (Figure 2a). The asymptotic and exact distributions must be highly similar for the asymptotic *p*-value *p*_asymptotic_ to be accurate. We measured goodness of fit of the Gaussian distribution to the exact null distribution by calculating the Kolmogorov–Smirnov (KS) statistic; this statistic ranges from zero to one, with smaller values indicating better fit of the Gaussian distribution to the exact null distribution. We reported the KS statistic for both example pairs in the panels of the plot.

Next, we calculated *p*_ratio_, defined as the ratio of the exact *p*-value *p*_exact_ to the asymptotic *p*-value *p*_asymptotic_, for each of the the 5,400 negative control pairs sampled from the Frangieh IFN-*γ* data. A *p*_ratio_ value of one indicates that the asymptotic and exact *p*-values coincide; a *p*_ratio_ value of greater than one (resp., less than one), on the other hand, indicates inflation (resp., deflation) of the asymptotic *p*-value relative to the exact *p*-value. We sought to explore visually how a small effective sample sizes lead to degradation of the Gaussian approximation, thereby resulting in *p*-value miscalibration (as reflected by *p*_ratio_ values that deviate from one). To this end, we plotted *p*_ratio_ versus goodness of fit of the the Gaussian distribution to the exact null distribution (as quantified by the KS statistic) for each pair (Figure 2b). We colored the points according to their effective sample size. Pairs 1 and 2 from Figure 2a were annotated in Figure 2b.

Finally, to directly assess the impact of sparsity on calibration, we applied Seurat-Wilcox to the IFN-*γ* negative control data, binning the pairs into five categories based on their effective sample size. The bins were defined by effective sample sizes in the ranges [7,10], [11,16], [17,27], [28,46], and [47,121]. The bins were constructed such that an approximately equal number of pairs would fall into each bin. We observed that as the effective sample size increased, the Seurat-Wilcox *p*-values converged to uniformity, illustrating that sparsity is a cause of the miscalibration of Seurat-Wilcox.

#### Confounding

We first explored how the variable of biological replicate confounded the Papalexi (gene modality) data. The Papalexi data were generated and sequenced across three independent experimental replicates, which we labeled “R1,” “R2,” and “R3”. (The original data contained a fourth biological replicate as well, but this replicate was removed by the original authors, as it was deemed to be of low quality.) We explored the relationship between biological replicate and a given NT gRNA (“NTg4”) and a given gene (*FTH1*). We plotted the fraction of cells in each biological replicate that received the NT gRNA (Figure 2d, left); additionally, we created a violin plot of the relative expression of the gene across biological replicate. (The relative expression *r*_*i*_ of the gene in cell *i* was defined as *r*_*i*_ = 1000 *·* log (*u*_*i*_*/l*_*i*_ + 1), where *u*_*i*_ was the UMI count of the gene in cell *i*, and *l*_*i*_ was the library size of cell *i*. The violin plots were truncated at a relative expression level of 50). We superimposed boxplots indicating the 25th, 50th, and 75th percentiles of the empirical relative expression distributions on top of the violin plots (Figure 2d, right). We observed clear visual evidence that biological replicate impacted both NTg4 and *FTH1*, creating a confounding effect.

Next, we extended the above analysis to investigate the entire set of NT gRNAs and genes. First, we tested for association between each NT gRNA and biological replicate. To this end, we constructed a contingency table of gRNA presences and absences across biological replicate, testing for significance of the contingency table using a using a Fisher exact test (as implemented in the R function `fisher.test`). Next, we tested for association between the relative expression of each gene and biological replicate. To do so, we fit two NB regression models to each gene; the first contained only library size as a covariate, while the second contained both library size *and* biological replicate as covariates. We compared these two models via a likelihood ratio test, yielding a *p*-value for the test of association between relative gene expression and biological replicate. Finally, we created QQ plots of the resulting *p*-values (Figure S6; gRNA *p*-values, left; gene *p*-values, right). An inflation of the *p*-values across modalities suggested that the bulk of gene-NT gRNA pairs was confounded by biological replicate.

Finally, we directly assessed the impact of adjusting for biological replicate (alongside other potential confounders) by applying two variants of NB regression to the Papalexi (gene modality) negative control data: (i) NB regression with library size (only) included as a covariate, and (ii) NB regression with library size as well as all potential confounders (including biological replicate) included as covariates. We plotted the negative control *p*-values on a QQ plot (Figure 2e). The variant of NB regression with confounders included as covariates exhibited superior calibration, demonstrating that confounding is an analysis challenge. To reduce the effect of sparsity (i.e., the first analysis challenge), we restricted our attention in this plot to gene-gRNA pairs with an effective sample size greater than 10.

#### Model misspecification

To explore the analysis challenge of model misspecification, we applied NB regression to the Frangieh IFN-*γ* negative control data. As in Figure 2c (in which we applied Seurat-Wilcox to the Frangieh IFN-*γ* negative control data), we partitioned the pairs into five categories based on the effective sample size of each pair. As the number of nonzero treatment cells increased, the NB regression *p*-values failed to converge to uniformity (in contrast to the Seurat-Wilcox *p*-values). The key difference between Seurat-Wilcox and NB regression is that the former is a nonparametric method while the latter is parametric method. Thus, we reasoned that miscalibration of the NB regression *p*-values likely was due to misspecification of the NB regression model. (We note that miscalibration of the NB regression *p*-values likely was not due to confounding, as Seurat-Wilcox, which does not adjust for confounding, was well-calibrated for pairs with high expression levels.)

### SCEPTRE (low-MOI) overview

Consider a given gene and perturbation. We call the cells that contain the targeting perturbation the “treatment cells” and those that contain an NT perturbation the “control cells.” Suppose there are *n* cells across treatment and control groups. Let *Y* = [*Y*_1_, …, *Y*_*n*_]^*T*^ be the vector of raw gene (or protein) expressions, and let *X* = [*X*_1_, …, *X*_*n*_]^*T*^ be the vector of perturbation indicators, where an entry of one (resp., zero) indicates presence of the targeting (resp. NT) perturbation. Finally, for cell *i* ∈ {1, …, *n*}, let *Z*_*i*_ be the *p*-dimensional vector of technical factors for cell *i* (containing library size, batch, etc.). We include an entry of one in each *Z*_*i*_ to serve as an intercept term. Let *Z* be the *n × p* matrix formed by concatenating the *Z*_*i*_s, and let [*X, Z*] be the *n ×* (*p* + 1) matrix formed by concatenating *X* and *Z*.

We model *Y*_*i*_ as a function of *X*_*i*_ and *Z*_*i*_ via an NB generalized linear model (GLM):

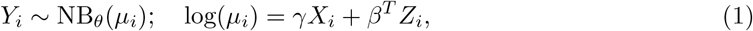

where NB_*θ*_(*μ*_*i*_) denotes a negative binomial distribution with mean *μ*_*i*_ and size parameter *θ*, and *γ* ∈ ℝ and *β* ∈ ℝ^*p*^ are unknown constants. (In fact, SCEPTRE in theory is compatible with arbitrary GLMs, including Poisson GLMs, which may be more appropriate for highly sparse data.)

SCEPTRE is a permutation test that uses as its test statistic the *z*-score that results from testing the hypothesis *γ* = 0 in the model (1). We present the basic SCEPTRE algorithm in Algorithm 1. Several key accelerations speed Algorithm 1 by multiple orders of magnitude.

#### Algorithm 1

Basic SCEPTRE algorithm.

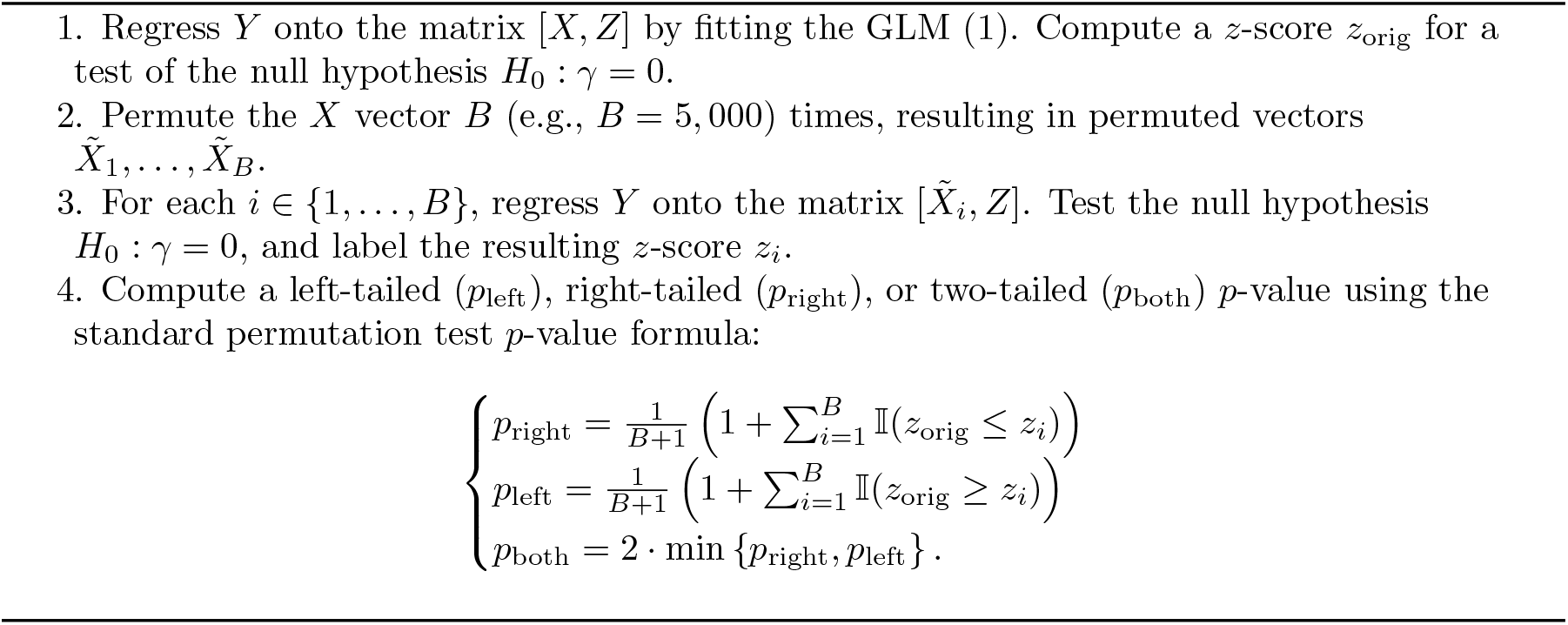

#### Acceleration 1: Score test

First, we use a GLM score test to compute the test statistics *z*_orig_, *z*_1_, …, *z*_*B*_. Consider the following simplified NB GLM in which the gene expression *Y*_*i*_ is modelled as a function of the technical factor vector *Z*_*i*_ only:

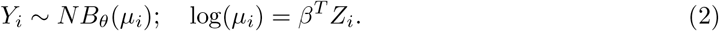

Regressing *Y* onto *Z* by fitting the GLM (2) produces estimates 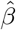 and 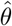 of the coefficient vector *β* and the size parameter *θ*, respectively, under the null hypothesis of no relationship between the gRNA indicator and the gene expression. Denote the *i*th fitted mean of the model by 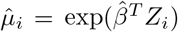, and let 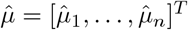 be the vector of fitted means. We can test the gRNA indicator vector *X* for inclusion in the fitted model by computing a score statistic *z*_score_, as follows.

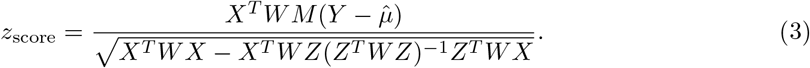

This expression is derived in the section “Derivation of the expression for the GLM score test statistic” of Additional file 1. Here, *W* and 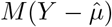 are a matrix and vector, respectively, that depend on the fitted means 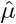, gene expressions *Y*, and estimated size 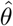:

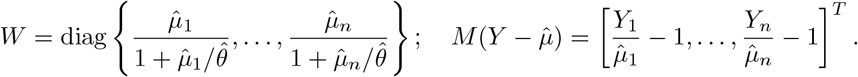

The vector 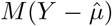 is a quantity called the working residual. The score statistic (3) is asymptotically equivalent to the Wald or likelihood ratio statistic that one obtains by testing *H*_0_ : *γ* = 0 in the full model (1). However, unlike the Wald statistic, the score statistic only depends on a fit of the model under the null hypothesis. SCEPTRE (with score statistic; Algorithm 2) exploits this useful property of the score statistic to accelerate the basic SCEPTRE algorithm.

##### Algorithm 2

SCEPTRE (with score statistic) algorithm.

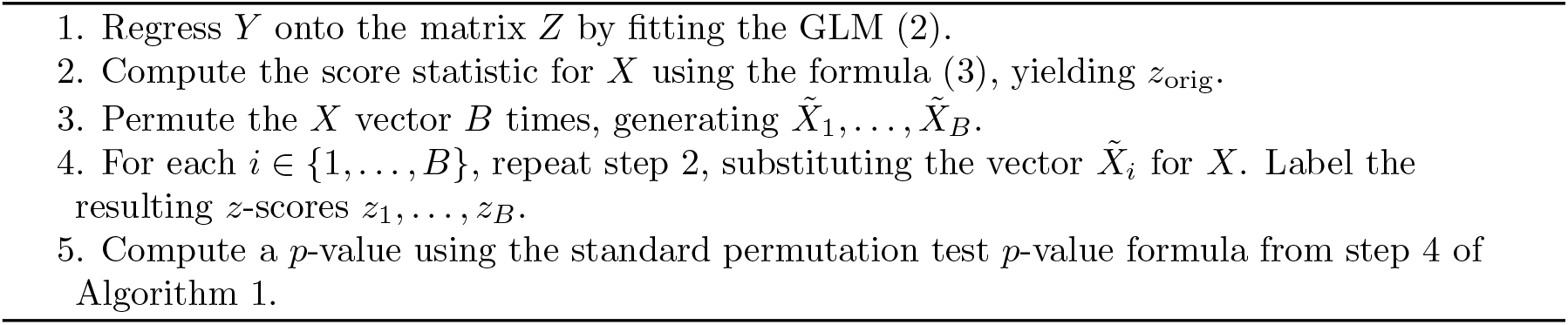

#### Acceleration 2: A fast score test for binary treatments

Calculating the score statistic (3) is not trivial. The quadratic form

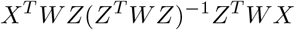

in the denominator of (3) is hard to compute, as the matrix *WZ*(*Z*^*T*^ *WZ*)^*−*1^*Z*^*T*^ *W* is a large, dense matrix. The classical solution is to algebraically manipulate the score statistic so that it can be evaluated via a QR decomposition. However, the QR decomposition approach does not leverage the structure in *X* when *X* contains many zeros (as is the case in single-cell CRISPR screen analysis). We therefore devised an alternate strategy for computing the score statistic that instead is based on a spectral decomposition; the proposed strategy is tens to hundreds of times faster than the QR decomposition approach when the treatment vector is sparse, as shown in section “Comparing the spectral decomposition algorithm to the QR decomposition algorithm for computing GLM score tests” of Additional file 1.

First, observe that *Z*^*T*^ *WZ* is a symmetric matrix. Thus, *Z*^*T*^ *WZ* can be spectrally decomposed as *Z*^*T*^ *WZ* = *U*^*T*^ Λ*U*, where *U* is an orthonormal matrix and Λ is a diagonal matrix of eigenvalues. Exploiting this decomposition, we can express the quadratic form in the denominator of (3) as follows:

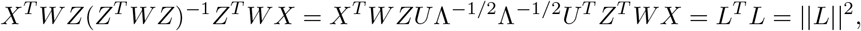

where *L* = Λ^*−*1*/*2^*U*^*T*^ *Z*^*T*^ *WX* is a *p*-dimensional vector. Evaluating the above expression reduces to computing the vector *L* and then summing over the squared entries of *L*, which is fast. This observation motivates Algorithm 3, which computes the score statistics for *X*, 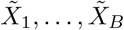 via a spectral decomposition.^3^ The inner product and matrix-vector multiplication operations of step 3 are extremely fast because *X*_curr_ is sparse. Furthermore, we program step 3 in C++ (via Rcpp[45]) for maximum speed.

##### Algorithm 3

Computing the GLM score statistics for *X*, 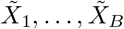 via spectral decomposition. Below, *w* is the *n*-dimensional vector constructed from the diagonal entries of *W*.

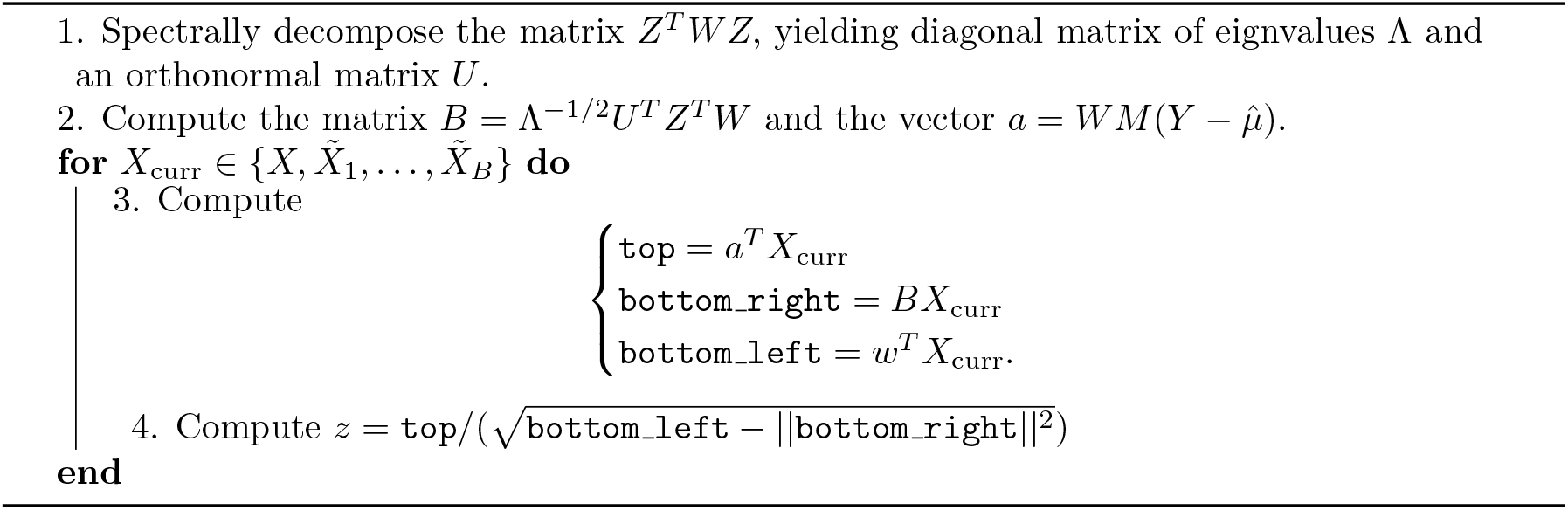

#### Acceleration 3: Adaptive permutation testing

Computing a large number of permutation resamples for a gene-gRNA pair that yields an unpromising *p*-value after only a few thousand resamples is wasteful. To reduce this inefficiency, we implement a two-step adaptive permutation testing scheme. First, we compute the *p*-value of a given gene-gRNA pair out to a small number (e.g., *B*_1_ = 500) of resamples. If this initial *p*-value is unpromising (i.e., if it exceeds some pre-selected threshold of *p*_thresh_, where *p*_thresh_ ≈ 0.01), then we return this *p*-value to the user. Otherwise, we draw a larger number (*B*_2_ = 5, 000) of fresh resamples and compute the *p*-value using this second set of resamples. As most pairs are expected to be null (and thus yield unpromising *p*-values), this procedure eliminates most of the compute associated with carrying out the permutation tests.

#### Acceleration 4: Skew-normal fit

The null distribution of the test statistics *z*_1_, …, *z*_*B*_ converges to a standard Gaussian distribution as the number of cells increases. Thus, to compute a precise *p*-value using a small number of permutations, we fit a skew-normal distribution to the set of null statistics. We then compute a *p*-value by evaluating the tail probability of the fitted skew-normal distribution at the observed test statistic *z*_obs_. If the skew-normal fit to the null statistics is poor (an event that happens rarely), we instead return the standard permutation test *p*-value. We fit the skew-normal distribution via a method of moments estimator and evaluate the skew-normal tail probability via the C++ Boost library. All operations involving the skew-normal distribution are fast.

#### Acceleration 5: Recycling compututation across permutation tests via IWOR sampling

When carrying out a permutation test to test for association between a gene expression vector *Y* = [*Y*_1_, …, *Y*_*n*_]^*T*^ and a perturbation indicator vector *X* = [*X*_1_, …, *X*_*n*_]^*T*^, SCEPTRE (low-MOI) randomly permutes the perturbation indicator vector *B* times, where *B* is some large number (e.g., *B* ≈ 5, 000). Unfortunately, randomly permuting the perturbation indicator vector *B* times is slow; this cost becomes prohibitive when testing many perturbation-gene pairs. We therefore derived a novel strategy for “sharing” a set of *B* randomly permuted indicator vectors across all perturbation-gene pairs, even pairs with different numbers of cells containing the targeting perturbation. This strategy — which we call “inductive without replacement” (IWOR) sampling — considerably reduces the cost associated with applying SCEPTRE to the data. (In fact, this method is generic, compatible with any permutation-based single-cell CRISPR screen association testing method.) IWOR sampling is described in section “Inductive without replacement sampling” of Additional file 1.

### Statistical robustness property of SCEPTRE

SCEPTRE empirically demonstrated a robustness property that we term “confounder adjustment via marginal permutations,” or “CAMP.” We observed evidence of CAMP across our simulation studies (Figure 4; Additional file 1: Figures S7, and S11) and real data analyses. We describe CAMP in greater detail here. For simplicity we consider the version of SCEPTRE that does *not* involve fitting a skew-normal distribution to the null test statistics and instead computes the standard permutation test *p*-value by directly comparing the observed test statistic to the null test statistics. If at least one of the following conditions holds, the left-, right-, and two-tailed SCEPTRE *p*-values are valid: (i) the perturbation is unconfounded (i.e., the vector of technical factors *Z*_*i*_ contains all possible confounders, and *Z*_*i*_ is independent of *X*_*i*_); (ii) the NB GLM (1) is correctly specified up to the size parameter *θ* and the effective sample size is sufficiently large. We reasoned that CAMP enabled SCEPTRE to address the core single-cell CRISPR screen analysis challenges of sparsity, confounding, and model misspecification both in theory and practice. Our evidence of CAMP is empirical; we intend to derive a mathematical proof of CAMP in a followup, more theoretical work.

### CAMP simulation study details

We conducted a simulation study (Additional file 1: Figure S7) to demonstrate the existence and utility of the CAMP phenomenon. We based the simulation study on a gene (namely, *CXCL10*) and perturbation (namely, “CUL3”) from the Papalexi data. Following the notation introduced in Section SCEPTRE (low-MOI) overview, let *Y* = [*Y*_1_, …, *Y*_*n*_]^*T*^ denote the vector of gene expressions of *CXCL10* and *X* = [*X*_1_, …, *X*_*n*_]^*T*^ the vector of perturbation indicators of “CUL3.” Next, let *Z*_*i*_ ∈ ℝ^*p*^ denote the vector of technical factors of the *i*th cell (for *i* ∈ {1, …, *n*}), and let *Z* denote the *n×p* matrix formed by assembling the *Z*_*i*_s into a matrix. We regressed *Y* onto *Z* by fitting the GLM (2), yielding estimates 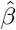 for *β* and *θ*^*\**^ for *θ* under the null hypothesis of no association between the perturbation and gene. An examination of 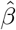 revealed that the gene expressions *Y* were moderately associated with the technical factors *Z*. Letting 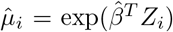 denote the fitted mean of cell *i*, we sampled *B* i.i.d. synthetic expressions 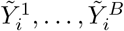 from an NB model with mean 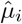 and size parameter *θ*^*\**^. We then constructed *B* synthetic gene expression vectors 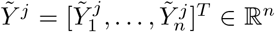 for *j* ∈ {1, …, *B*}. Next, we generated a synthetic perturbation indicator vector 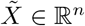 such that 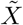 was independent of *Z*. To this end, we marginally sampled synthetic perturbation indicators 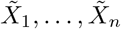 i.i.d. from a Bernoulli model with mean 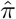, where 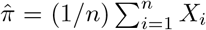 was the fraction of cells that received the targeting perturbation. (The observed perturbation indicator vector *X* was moderately associated with *Z*.)

We assessed three methods in the simulation study: NB regression, SCEPTRE, and the standard permutation test. We deployed NB regression and SCEPTRE in a slightly different way than usual: we set the NB size parameter *θ* upon which these methods rely to a fixed value. (Typically, NB regression and SCEPTRE estimate *θ* using the data.) This enabled us to assess the impact of misspecification of the size parameter on the calibration of NB regression and SCEPTRE. We set the test statistic of the standard permutation test to the sum of the gene expressions in the treatment cells. We then generated *B* confounded (resp., unconfounded) datasets by pairing the synthetic response vectors 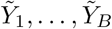 to the design matrix [*X, Z*] (resp., 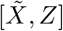). We applied the methods to the datasets twice: once setting the SCEPTRE/NB regression size parameter to the correct value of *θ*^*\**^, and once setting this parameter to the incorrect value of 5 *· θ*^*\**^. We displayed the results produced by the methods in each of the four settings (i.e., confounded versus unconfounded, correct versus incorrect specification of the size parameter; Additional file 1: Figure S7) on a QQ plot. We sought to show that SCEPTRE maintains calibration in all settings, while the standard permutation test and NB regression break down under confounding and incorrect specification of the size parameter, respectively.

### Positive control analysis

We grouped together gRNAs that targeted the same genomic location, referring to these grouped gRNAs as “gRNA groups”[5]. We constructed positive control pairs by coupling a given gRNA group to the gene or protein that the gRNA group targeted. We developed a Nextflow pipeline to apply all methods to analyze the positive control pairs of all datasets.

### ChIP-seq enrichment analysis

We obtained ChIP-seq data for CD14+ monocyte cultures with MCSF (10ng/ml) and stimulated with IFN-gamma (100U/ml) for 24 hours.[30] Peaks were screened based on their score, with the top 25% selected using the enrichment score as the criterion. We identified downstream target genes for transcription factors by identifying genes with transcription start sites located within 5kb upstream or downstream of a peak. Using ChIP-seq as the benchmark, our objective was to assess the consistency between the genes associated with a given transcription factor as identified by SCEPTRE and the downstream target genes determined by ChIP-seq. To achieve this, we calculated odds ratios and their corresponding p-values using a Fisher exact test on the contingency table comprised of genes found to be affected by knockout of the perturbed gene, as identified by SCEPTRE, and genes whose promoter regions overlapped with a ChIP-seq peak. We conducted this analysis for both STAT1 and IRF1. We also performed this analysis across competing methods, as shown in Additional file 1: Figure S9.

### Comparison of SCEPTRE (low-MOI) and SCEPTRE (high-MOI)

SCEPTRE (low-MOI) is a substantial statistical and computational extension of SCEPTRE (high-MOI). Below, we outline the ways in which SCEPTRE (low-MOI) differs from SCEPTRE (high-MOI) version 0.0.2, which is the version of SCEPTRE (high-MOI) available on our website at the time of submission.

- SCEPTRE (low-MOI) carries out inference via a permutation test, while SCEPTRE (high-MOI) does so via a conditional randomization test. Given that the low-MOI problem suffers from stronger sparsity, while the high-MOI problem suffers from greater confounding, we reasoned that permutations would yield better calibration in the low-MOI setting.
- SCEPTRE (low-MOI) uses a full GLM score statistic as its test statistic, while SCEPTRE (high-MOI) uses a distilled GLM score statistic. The full score statistic is more powerful than its distilled counterpart, yielding a greater number of discoveries. Moreover, the full score statistic used by SCEPTRE (low-MOI) is supported by a novel and fast algorithm for computing GLM score tests.
- SCEPTRE (low-MOI) leverages a novel algorithm for recycling computation across a large number of permutation tests, thereby considerably decreasing computational cost. This approach — which we term “inductive without replacement sampling” — is described in Section Inductive without replacement sampling.
- SCEPTRE (low-MOI) fits a skew-normal distribution to the null test statistics, while SCEPTRE (high-MOI) fits a skew-*t* distribution to the null test statistics. The skew-normal distribution admits a fast and numerically stable method-of-moments estimator, while the skew-*t* distribution requires a slow and (relatively) numerically unstable maximum likelihood estimator.
- SCEPTRE (low-MOI) checks for goodness of fit of the fitted skew-normal distribution before it uses the fitted distribution to compute a *p*-value. SCEPTRE (high-MOI), by contrast, does not check for goodness of fit of the fitted skew-*t* distribution, which potentially can lead to miscalibrated *p*-values for perturbation-gene pairs whose resampling distributions are not approximately skew-*t*-distributed.
- SCEPTRE (low-MOI) uses an adaptive permutation testing scheme to reduce the number of permutations computed for pairs with unpromising *p*-values. SCEPTRE (high-MOI), by contrast, does not leverage any sort of adaptive resampling scheme.
- SCEPTRE (high-MOI) is programmed entirely in R. SCEPTRE (low-MOI) by contrast, is programmed in a mix of C++ and R, with the computationally intensive portions programmed in C++.
- SCEPTRE (low-MOI) uses a considerably faster method than SCEPTRE (high-MOI) for fitting the negative binomial regression models.
- SCEPTRE (low-MOI) includes support for “pairwise quality control,” in which low-quality perturbation-gene pairs (defined as pairs whose effective sample size falls below some threshold) are detected and removed.
- SCEPTRE (low-MOI) can automatically construct negative control pairs that are “matched” to the discovery pairs in several respects; these negative control pairs can be used to assess the calibration of SCEPTRE (low-MOI) on a user-inputted dataset. SCEPTRE (high-MOI) does not have such functionality.
- SCEPTRE (low-MOI) by default uses the set of negative control cells as its control group; this choice is especially appropriate for gene-targeting screens. SCEPTRE (high-MOI), by contrast, uses the complement set as its control group, as this is the only option, since few (if any) cells contain exclusively non-targeting perturbations in the high-MOI setting.
- SCEPTRE (low-MOI) includes new functions for visualizing the results, including a function to create a volcano plot and a function to create a QQ plot with the discovery *p*-values superimposed on top of the negative control *p*-values.

Taken together, these extensions make SCEPTRE (low-MOI) faster, more memory efficient, more statistically powerful, more statistically robust, more numerically stable, and more user-friendly than SCEPTRE (high-MOI). We anticipate that many of these extensions can be applied to improve the high-MOI functionality of SCEPTRE as well. (Important differences between the two modules, however, including the choice of control group, will remain.) We intend to explore this possibility in future work.

### Methods not included in the benchmarking analysis

Several methods that recently have been proposed for single-cell CRISPR screen analysis were not included in our benchmarking study. First, guided sparse factor analysis (GSFA; introduced by Zhou et al.[39]) couples factor analysis to differential expression analysis to infer the effects of perturbations on gene modules and individual genes. GSFA is a Bayesian method, returning a posterior inclusion probability instead of a *p*-value for each test of association. Given that the methods that we studied in this work were frequentist (and thus returned a *p*-value), we deprioritized GSFA for benchmarking. Next, Normalisr (proposed by Wang[17]) is a method for single-cell differential expression, co-expression, and CRISPR screen analysis. Normalisr non-linearly transforms the gene expression counts to Gaussianity and then models the transformed counts via a linear model. We were unable to locate an example low-MOI single-cell CRISPR screen analysis in the Normalisr Github repository (although gene co-expression, case-control differential expression, and high-MOI CRISPR screen examples are available). Given this, and given the complexity of the Normalisr code-base, we deprioritized Normalisr for benchmarking. Finally, scMaGECK (proposed by Yang[14]) tests for association between CRISPR perturbations and gene expressions using a permutation test with a linear regression coefficient test statistic. Given that we carefully evaluated the high-MOI version of scMaGECK in our prior work[13], and given that MIMOSCA — benchmarked in this work — also is based on a permutation test with a linear regression coefficient test statistic, we deprioritized scMaGECK for benchmarking.

## Declarations

### Availability of data and materials

The code for this paper is contained across nine Github repositories. Navigate to github.com/Katsevich-Lab/sceptre2-manuscript (i.e., the second Github repository of those listed) for instructions on reproducing the analyses reported in this manuscript.

1. The `sceptre` package implements the SCEPTRE method. It is released under a GPL-3.0 license. The repository contains detailed tutorials and examples. katsevich-lab.github.io/sceptre
2. The `sceptre2-manuscript` repository [46, 47] contains code to reproduce all analyses reported in this paper. It is the main reproduction repository associated with this manuscript. It is released under a GPL-3.0 license. It is deposited at Zenodo with DOI 10.5281/zen-odo.10976334. github.com/Katsevich-Lab/sceptre2-manuscript
3. The `lowmoi` package implements the existing single-cell CRISPR screen analysis methods. (Methods originally written in Python are implemented via `reticulate`). github.com/Katsevich-Lab/lowmoi
4. The `undercover-grna-pipeline` repository contains the Nextflow pipeline to carry out negative control benchmarking analysis. github.com/Katsevich-Lab/undercover-grna-pipeline
5. The `pc-grna-pipeline` repository contains the Nextflow pipeline to carry out the positive control benchmarking analysis. github.com/Katsevich-Lab/pc-grna-pipeline
6. The `ondisc` package implements data structures that we use to store the single-cell expression data. github.com/timothy-barry/ondisc
7. The `import-frangieh-2021` repository imports and processes the Frangieh data. github.com/Katsevich-Lab/import-frangieh-2021
8. The `import-papalexi-2021` repository imports and processes the Papalexi data. github.com/Katsevich-Lab/import-papalexi-2021
9. The `import-schraivogel-2020` repository imports and processes the Schraivogel data. github.com/Katsevich-Lab/import-schraivogel-2020

Next, the results are available on DropBox (https://www.dropbox.com/scl/fo/5uw77telw7uvgatkj0xod/AKAPdd5jdrswtRXahVxY0Tk?rlkey=2mw1kzu4erztqzr9qd6f2ojxq&dl=0). Finally, the processed single-cell CRISPR screen data (stored in `ondisc v1.1.0` format) are available on Zenodo (DOI: 10.5281/zenodo.10976417) and on Dropbox (www.dropbox.com/sh/jekmk1v4mr4kj3b/AAAhznGqk-TIZKhW40xiU6ORa?dl=0).

## Competing interests

The authors declare that they have no competing interests.

## Funding

This study was supported by Analytics at Wharton, NSF grants DMS 2113072 and DMS 2310654, and NIH grant R01MH123184.

## Ethical Approval

Not applicable for this study.

## Author contributions

EK identified the research problem. TB, KM, and EK performed the analyses. TB and EK developed the method. TB implemented the low-MOI SCEPTRE software. KR and EK supervised the project. TB and EK wrote the manuscript with assistance from KM and KR.

## Acknowledgements

We thank Hugh MacMullan and Gavin Burris for extensive support in using the Wharton high performance computing cluster (HPCC). We thank Sophia Lu for preliminary work on benchmarking existing methodologies. We thank Ziang Niu for carrying out analyses related to modeling the SCEPTRE resampling distributions. We thank John Morris for help in designing the computational experiments and providing feedback on an early draft. We thank Chris Frangieh for providing instructions on downloading and processing the Frangieh data and using the MIMOSCA method. We thank Rahul Satija for clarifying several points about the Papalexi data. We thank Stephanie Hicks, Kasper Hansen, and the Hicks and Hansen Labs at Johns Hopkins University for detailed feedback and recommendations about additional analyses to conduct. We thank Luca Pinello, Mike Love, and the CRISPR Working Group at the Impact of Genomic Variation on Function (IGVF) Consortium for helpful comments and discussion. Finally, we thank two anonymous reviewers for comments that considerably improved the manuscript.

## Additional files

Additional file 1 contains the supplementary tables and figures referenced in the main text, additional mathematical details of SCEPTRE (low-MOI), and several additional empirical analyses.

## Additional file 1 for

### Organization of Additional file 1

This document contains supplementary materials for the paper *Robust differential expression testing for single-cell CRISPR screens at low multiplicity of infection*. First, we present the supplementary tables (i.e., Tables S1 — S3) and figures (i.e., S1 — S8) referenced in the main text. Next, we provide additional mathematical details about the SCEPTRE (low-MOI) method, including a discussion of inductive without replacement (IWOR) sampling. Finally, we present several additional empirical analyses, including a comparison of the score statistic and the difference-in-residual-means statistic in the context of permutation testing, a comparison of the spectral decomposition algorithm and the QR decomposition algorithm for computing GLM score tests, and an analysis of the number of cells required per perturbation to ensure adequate power.

## S1 Supplementary tables and figures referenced in the main text

**Table S1:** Datasets analyzed in this work. The first column indicates the name of a low-MOI single-cell CRISPR screen paper; the second column indicates the datasets that we obtained from that paper; and the subsequent columns indicate the (paper-specific) biological attributes of the data, including CRISPR modality, technology platform, target type, cellular modality measured, and cell type. ^*\**^The Frangieh data also contain protein measurements, but we focus exclusively on the gene modality in this work.

| Paper | Datasets | CRISPR modality | Platform | Target | Modality measured | Cell type |
| --- | --- | --- | --- | --- | --- | --- |
| Frangieh 2021 | co-culture, control, IFN-γ (3) | CRISPRko | Perturb-CITE seq | Gene TSSs | Gene expressions* | TIL |
| Papalexi 2021 | ECCITE screen (1) | CRISPRko | ECCITE-seq | Gene TSSs | Gene and protein expressions | THP1 |
| Schraivogel 2020 | Enhancer screen (1) | CRISPRi | Targeted perturb-seq | Enhancers | Gene expressions | K562 |
| - | Simulated dataset (1) | - | - | Gene TSSs | Gene expressions | - |

**Table S2:** Statistical attributes of the datasets. The number of genes, cells, targeting gRNAs, NT gRNAs, negative control pairs, and positive control pairs for each dataset. Neg., negative; pos., positive. *Schraivogel separately assayed two chromosomes: Chr11 and Chr8. Given the similarity of these assays, we pool together the negative and positive control pairs across assays.

| Dataset | N genes<br>(or proteins) | N cells | N targeting<br>gRNAs | N NT<br>gRNAs | N neg.<br>control<br>pairs | N pos.<br>control<br>pairs |
| --- | --- | --- | --- | --- | --- | --- |
| Frangieh Co-culture | 14,438 | 46,427 | 744 | 74 | 596,344 | 181 |
| Frangieh control | 15,449 | 30,486 | 744 | 74 | 528,239 | 170 |
| Frangieh IFN-γ | 14,654 | 50,053 | 744 | 74 | 565,502 | 181 |
| Papalexi (gene) | 14,559 | 20,729 | 101 | 9 | 100,458 | 25 |
| Papalexi (protein) | 4 | 20,729 | 101 | 9 | 36 | 2 |
| Schraivogel* | 82 (Chr11),<br>71 (Chr8) | 99,884 (Chr11),<br>88,715 (Chr8) | 3,073 (Chr11),<br>4,089 (Chr8) | 30 (Chr11),<br>30 (Chr8) | 4,693 (pooled) | 25 (pooled) |
| Simulated | 4439 | 10,000 | - | 25 | 108,510 | - |

**Table S3:** Analysis challenges addressed by each method. Each cell indicates whether the method in the row addresses the analysis challenge in the column. SCEPTRE (bottom row) is the only method that addresses all three analysis challenges. Note: MIMOSCA is excluded from this table, as we could not determine which analysis challenge(s) MIMOSCA addresses.

| Method | Sparsity | Confounding<br>(beyond<br>library size) | Model<br>misspecification |
| --- | --- | --- | --- |
| Seurat-Wilcox | No | No | Yes |
| Seurat-NB | No | No | No |
| *t*-test | Yes | No | No |
| MAST | No | No | No |
| KS test | No | No | Yes |
| NB regression (with covariates) | No | Yes | No |
| Standard permutation test | Yes | No | Yes |
| **SCEPTRE** | **Yes** | **Yes** | **Yes** |

**Table S4:** Benchmark comparison of the spectral decomposition algorithm and the QR decomposition algorithm for computing GLM score tests. Left column, sparsity level of the treatment vector (i.e, mean fraction of entries within the treatment vector equal to zero). Middle two columns, mean running time of the spectral decomposition method and QR decomposition method (95% confidence intervals in parentheses). Right column, “speedup” (i.e., the QR decomposition running time divided by the spectral decomposition running time).

| Sparsity | Spectral decomp. running time | QR decomp. running time | Speedup |
| --- | --- | --- | --- |
| 99.9% | 0.0057s (0.0056, 0.0058) | 1.55s (1.55, 1.56) | 272 |
| 99% | 0.0088s (0.0087, 0.0089) | 1.60s (1.60, 1.60) | 181 |
| 95% | 0.023s (0.023, 0.023) | 1.62s (1.61, 1.62) | 71 |
| 90% | 0.040s (0.040, 0.040) | 1.61s (1.61, 1.61) | 40 |
| 50% | 0.147s (0.146, 0.147) | 1.62s (1.62, 1.62) | 11 |

**Figure S1:**
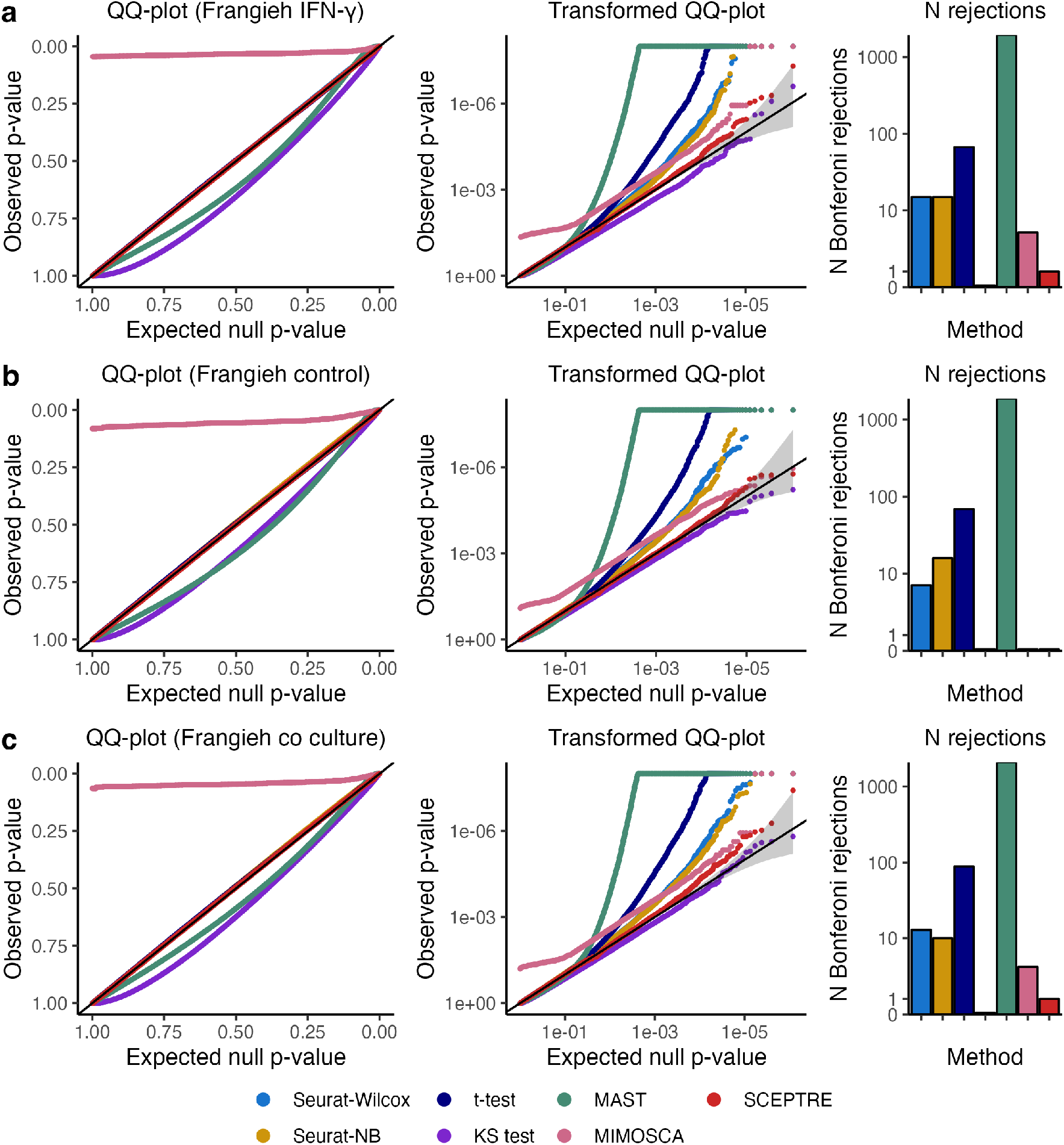
Calibration results for all methods on Frangieh IFN-*γ*, Frangieh control, and Frangieh co-culture negative control data. Left, untransformed QQ plots; middle, negative log-10 transformed QQ plots; right; number of false rejections after a Bonferroni correction at level 0.1.

**Figure S2:**
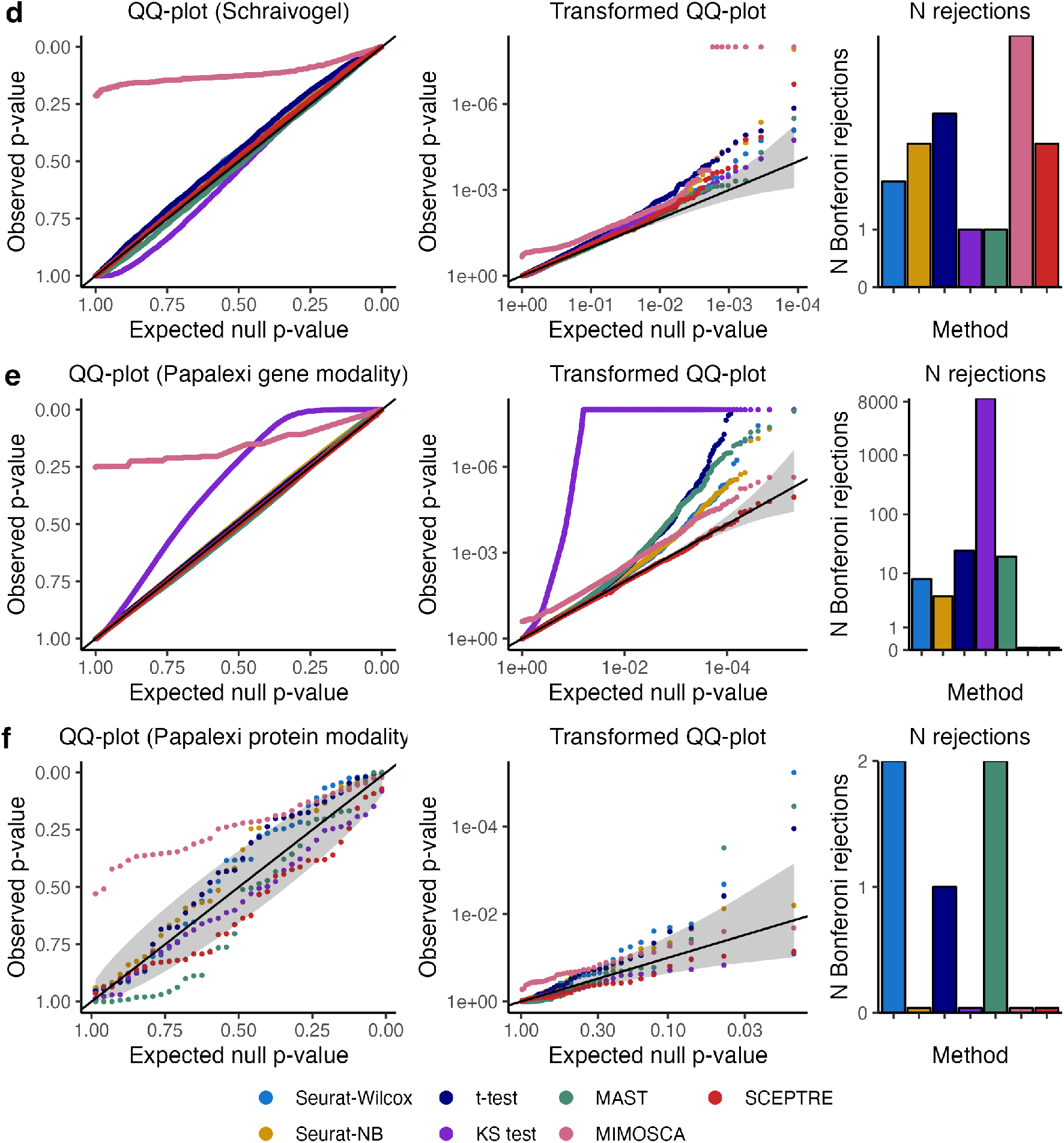
Calibration results for all methods on Schraivogel, Papalexi (gene modality), and Papalexi (protein modality) negative control data. Interpretation is the same as in Figure S1.

**Figure S3:**
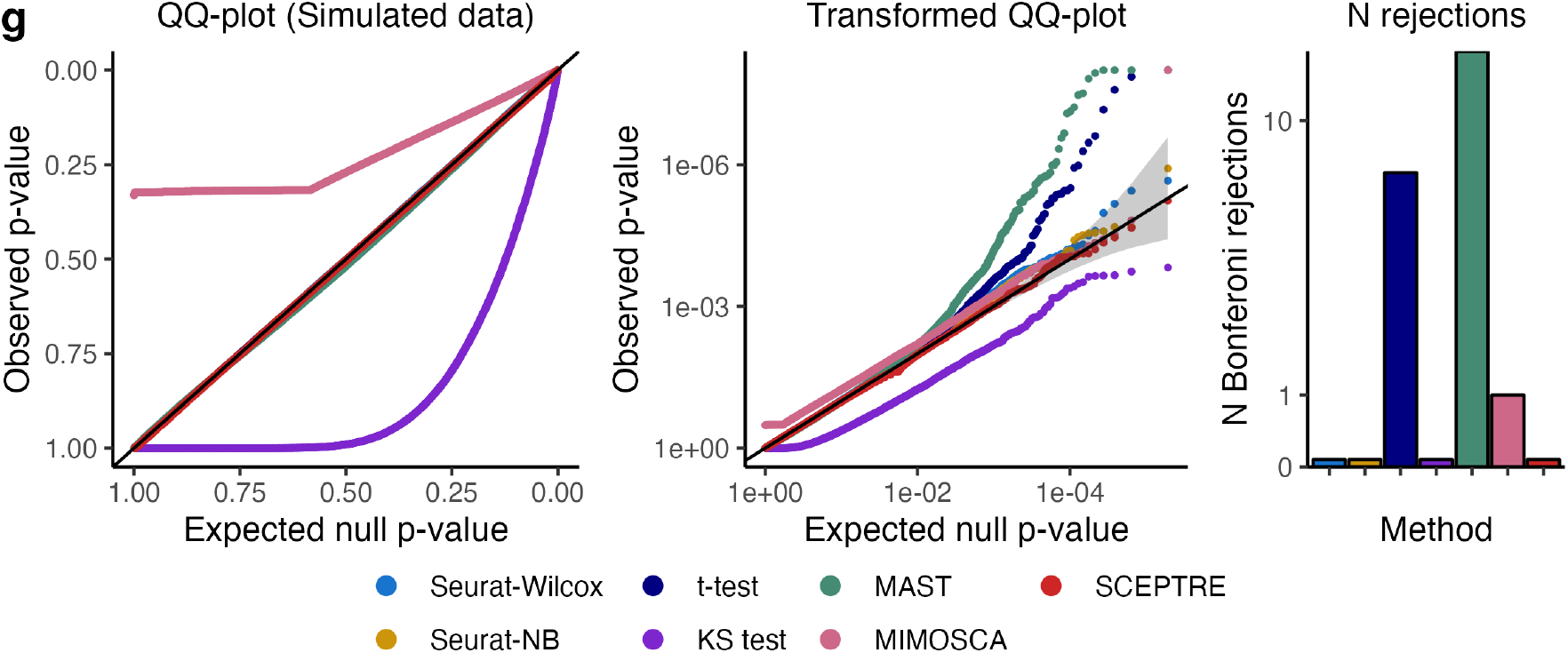
Calibration results for all methods on simulated data. Interpretation is the same as in Figure S1.

**Figure S4:**
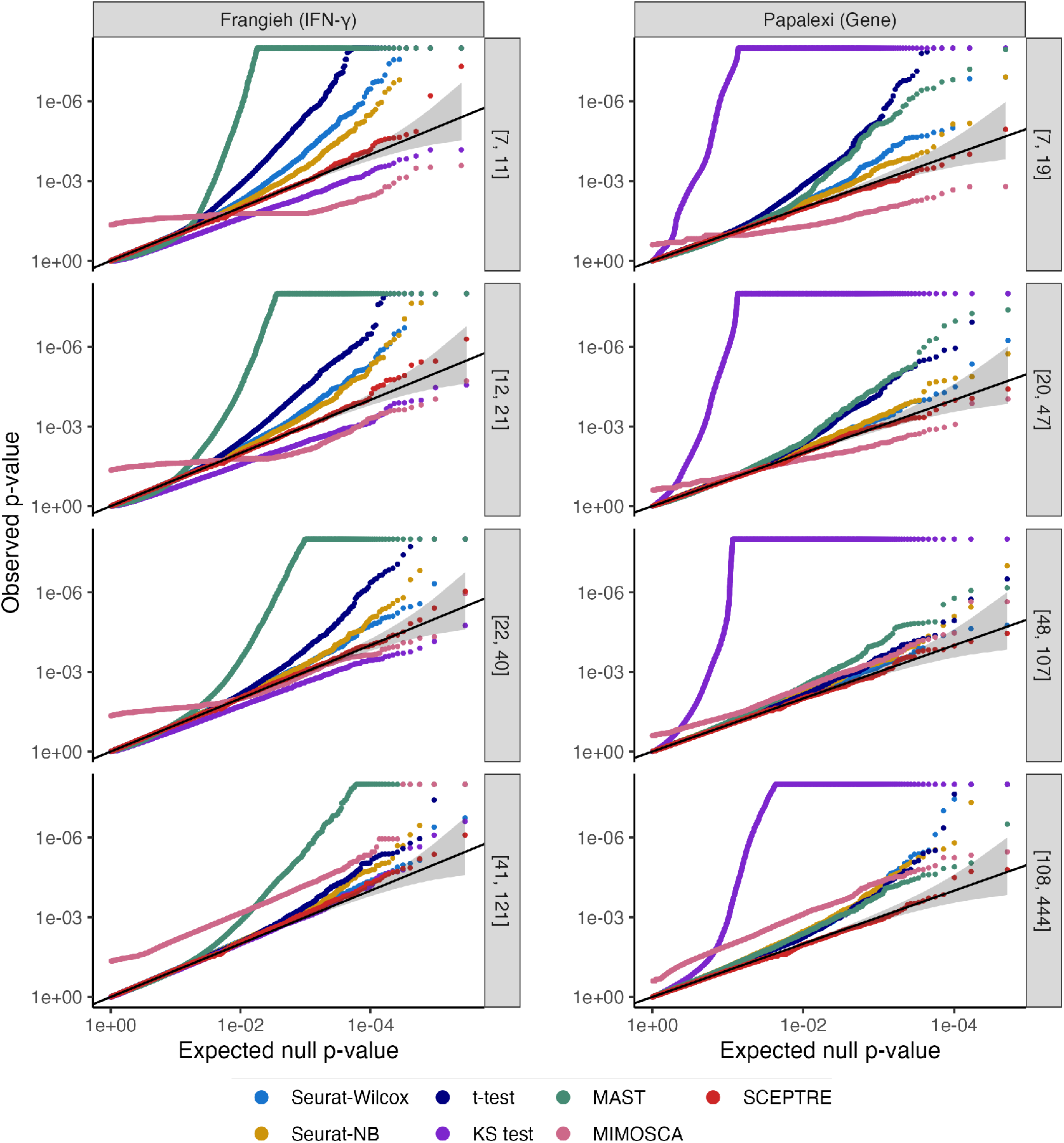
Calibration results for all methods on Frangieh (IFN-*γ*) and Papalexi (gene modality) datasets, stratified by effective sample size. Negative control gene-gRNA pairs are partitioned into four bins of approximately equal size based on the number of treatment cells with nonzero expression in a given pair. The interval on the right-hand side of each panel indicates the minimum and maximum number of treatment cells with nonzero gene expression for pairs in that bin. Some methods (e.g., Seurat-Wilcox on the Frangieh IFN-*γ* data) exhibit better calibration as the number of treatment cells with nonzero expression increases (i.e., as sparsity decreases).

**Figure S5:**
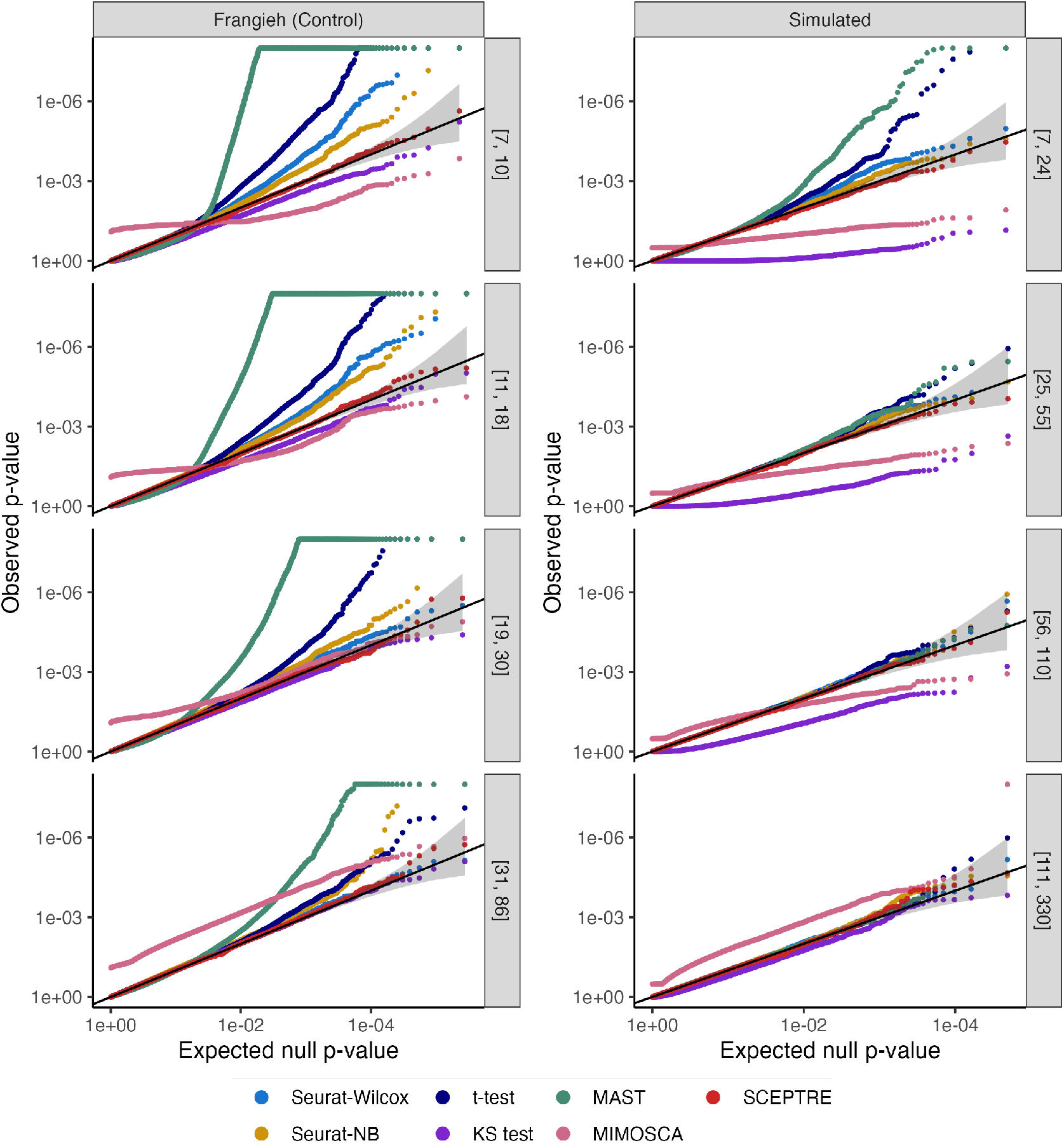
Calibration results for all methods on Frangieh (control) and simulated datasets, stratified by effective sample size. This figure is identical to Figure S4 but displays results on the Frangieh (control) and simulated datasets.

**Figure S6:**
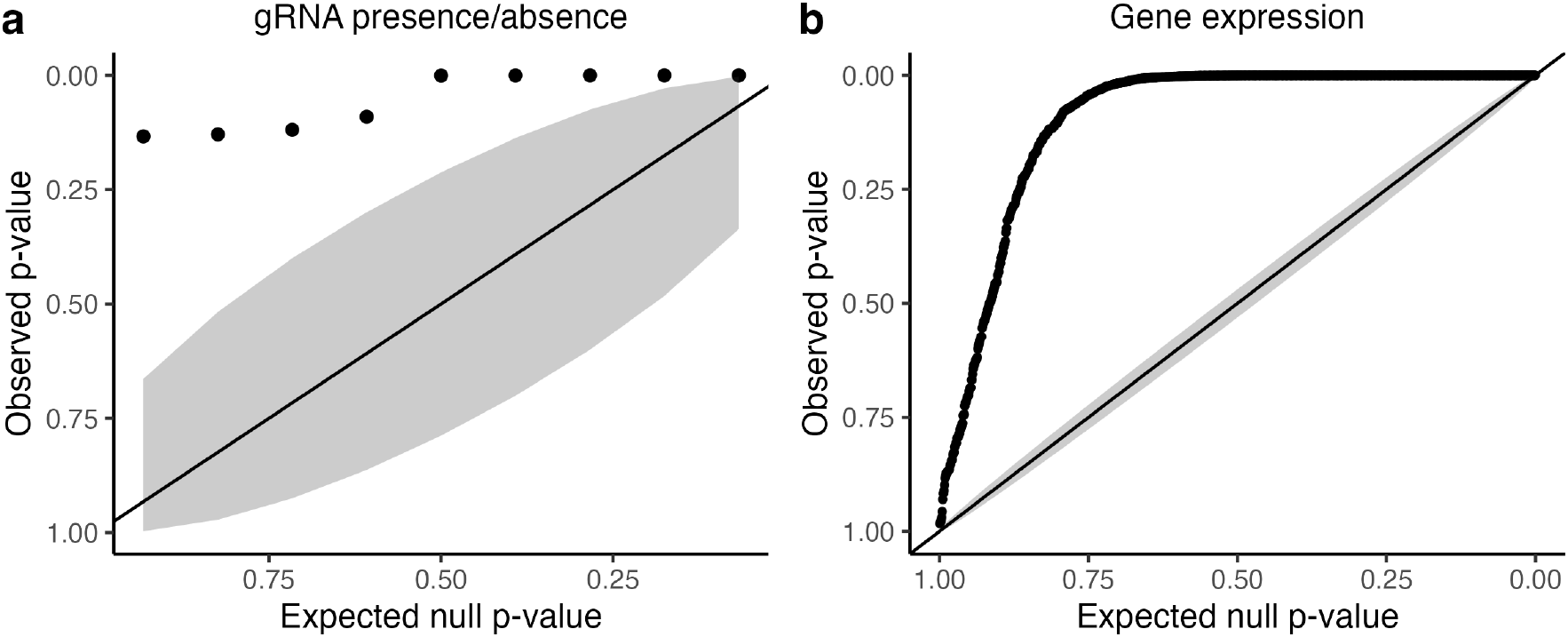
Confounding due to biological replicate on the Papalexi (gene modality) data. We conducted a test of association between each NT gRNA and biological replicate. We plotted the resulting *p*-values on a QQ plot (left); each point in the plot corresponds to a different NT gRNA. Likewise, we carried out a test of association between the expression of each gene and biological replicate (right); each point in the plot similarly corresponds to a different gene. We observed that the two sets of *p*-values were inflated, indicating that the bulk of the NT gRNAs and the bulk of the genes was associated with biological replicate. Because biological replicate affected *both* gRNA presence/absence *and* gene expression, we reasoned that biological replicate acted as a confounder on the Papalexi data. The section Details of the investigation into the core analysis challenges (“Confounding” subsection) describes how the tests of association were conducted.

**Figure S7:**
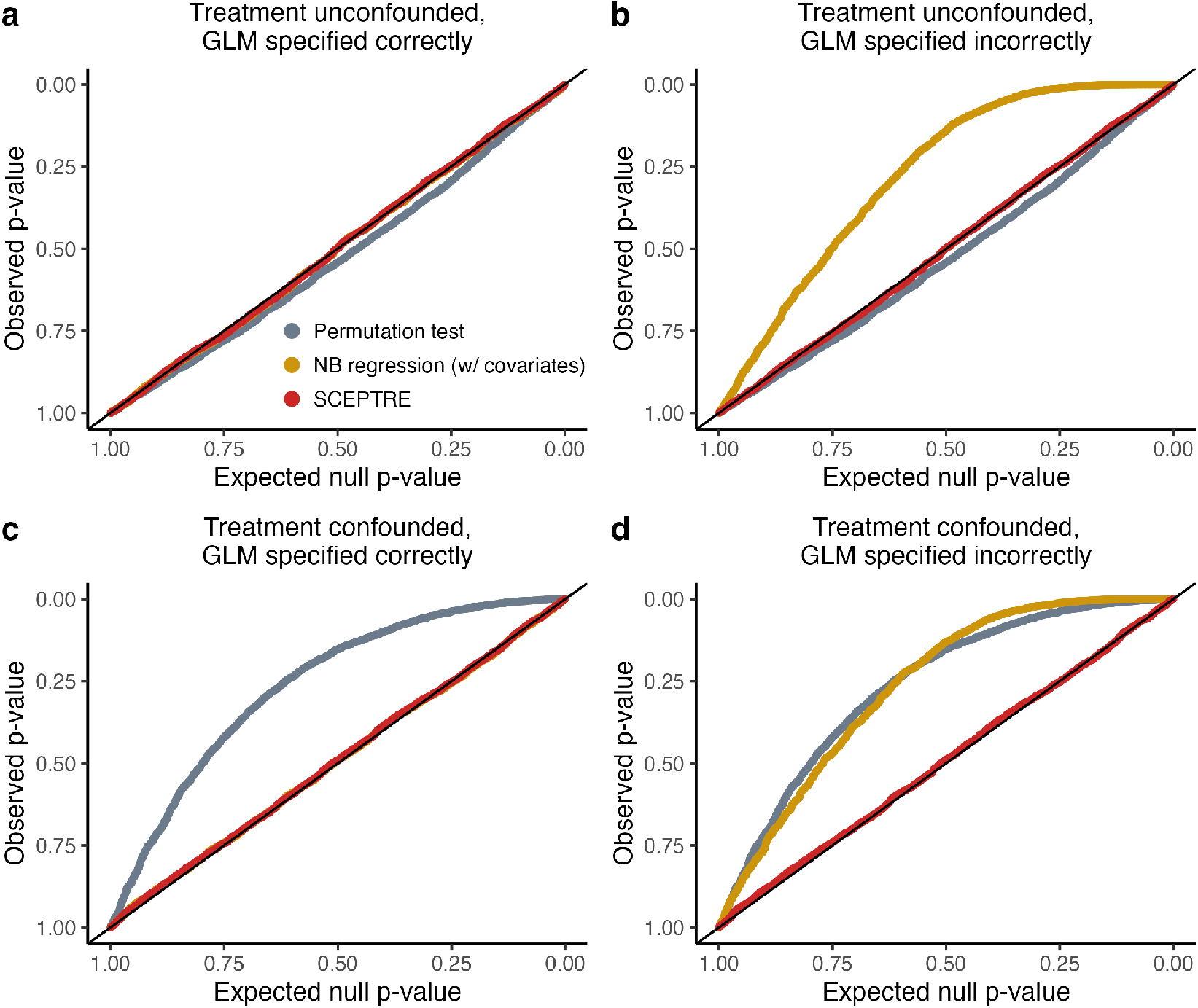
Demonstration of the CAMP (“confounder adjustment via marginal permutations”) phenomenon on realistic semi-synthetic data. Application of a standard permutation test, NB regression, and SCEPTRE to realistic semi-synthetic data generated under two conditions: confounded and unconfounded. Panels **a** and **b** (resp., **c** and **d**) show the results on the unconfounded (resp., confounded) data; meanwhile, panels **a** and **c** (resp., **b** and **d**) show the results under correct (resp. incorrect) specification of the negative binomial size parameter. The permutation test (gray) works well when the data are unconfounded (panels **a** and **b**) but breaks down in the presence of confounding (panels **c** and **d**). On the other hand, NB regression is well-calibrated when the size parameter is correctly specified (panels **a** and **c**) but fails when the size parameter is misspecified (panels **b** and **d**). SCEPTRE is well-calibrated in all settings. We note that SCEPTRE is expected to break down when the (i) problem is confounded and the NB regression model is arbitrarily misspecified or (ii) the problem is confounded and the sparsity is high. Details of the simulation study are given in Section CAMP simulation study details.

**Figure S8:**
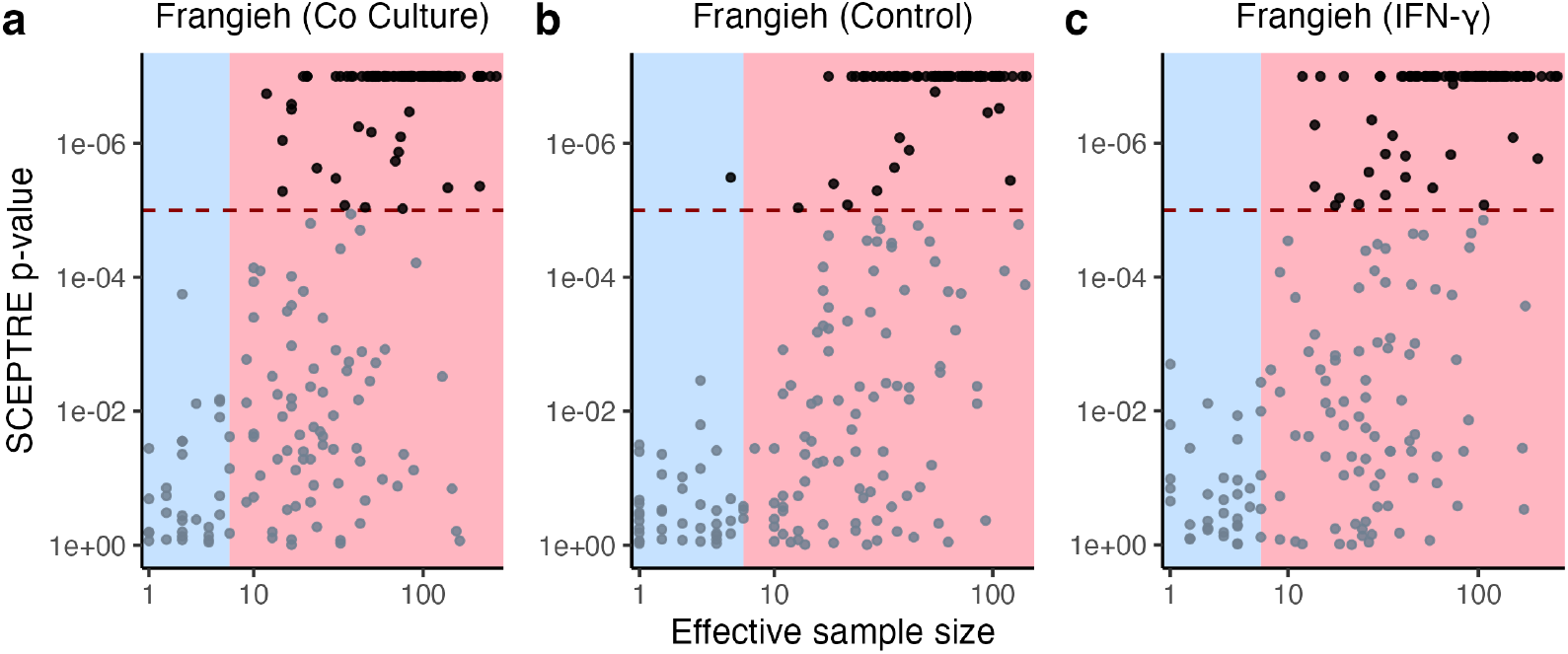
SCEPTRE’s power to detect associations increases as effective sample size increases. **a-c**, SCEPTRE *p*-value (truncated at 10^*−*6^) versus effective sample size for each pair on the Frangieh co-culture (**a**), control (**b**), and IFN-*γ* (**c**) positive control data. The horizontal dashed line is drawn at 10^*−*5^, demarcating a highly significant discovery. SCEPTRE makes only one rejection at a highly significant level on pairs for which the effective sample size less than seven (blue region).

**Figure S9:**
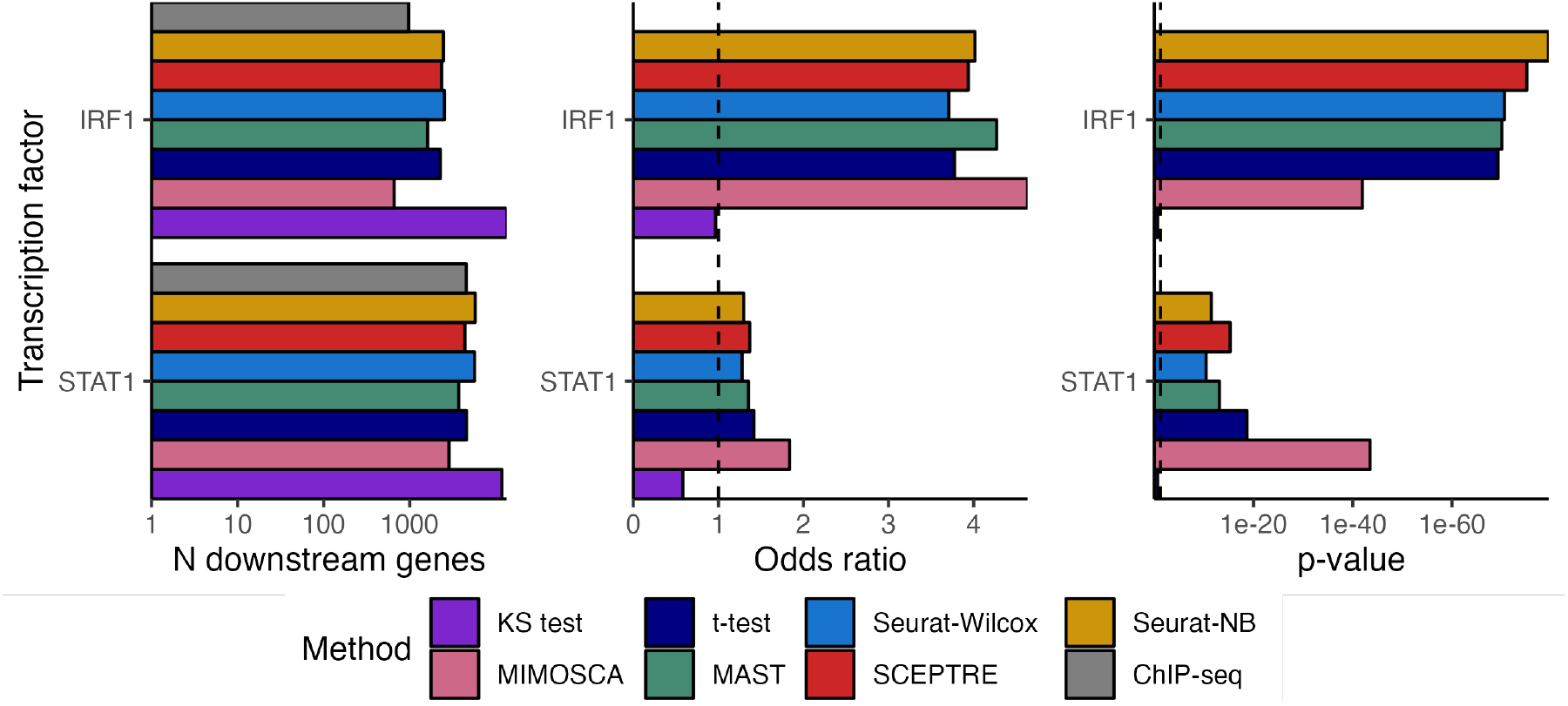
Comparison of methods for enrichment of ChIP-seq signal on the Papalexi data. Number of discoveries (after a BH correction at level 0.1), ChIP-seq enrichment odds ratio, and ChIP-seq enrichment p-value of each method for the transcription factors IRF1 and STAT1. The p-value quantifies the extent to which the discovery set of a given method exhibited enrichment for cell-type-relevant ChIP-seq signal. See section ChIP-seq enrichment analysis for a description of how the ChIP-seq data were used to identify the target genes of STAT1 and IRF1.

**Figure S10:**
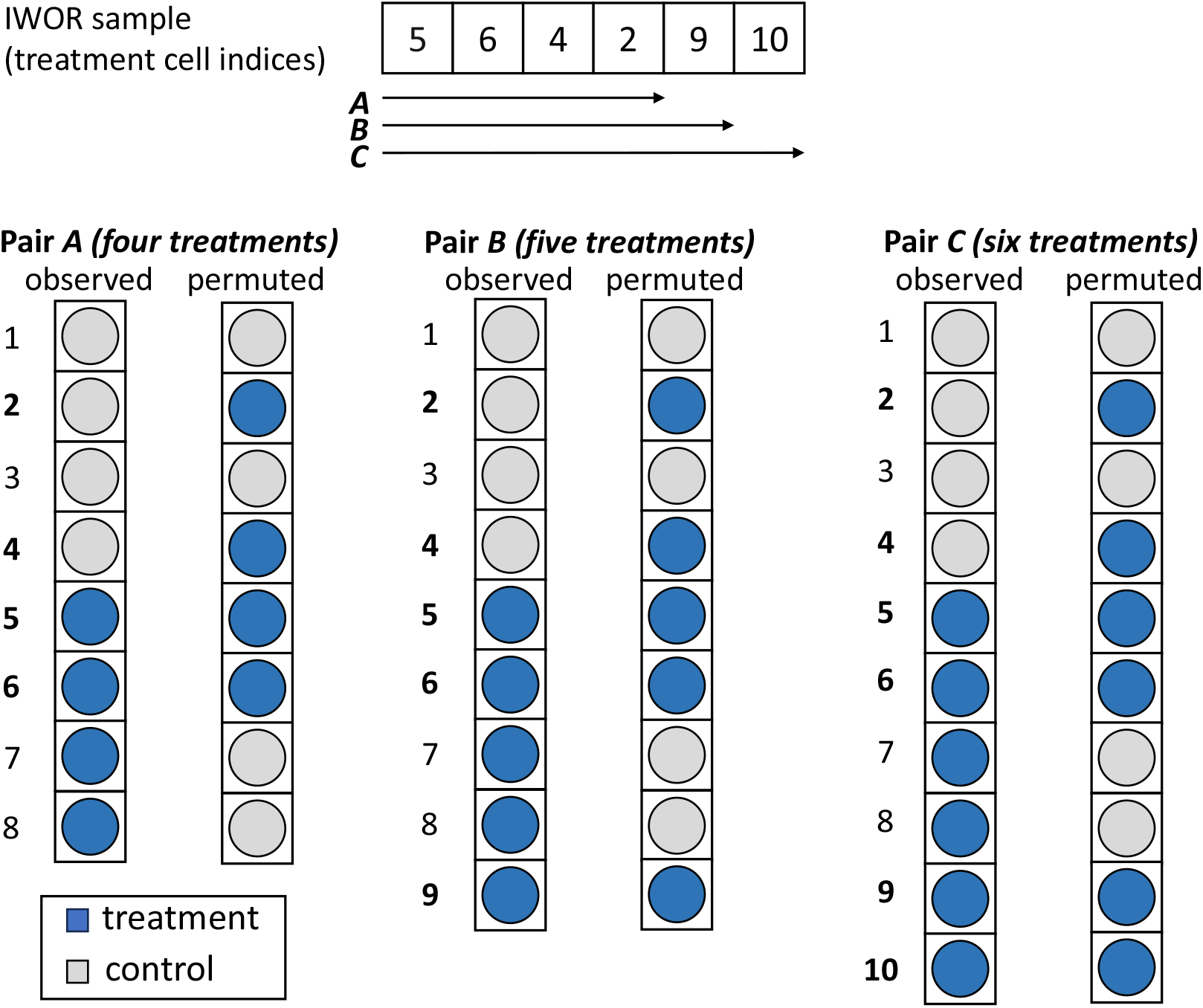
Inductive without replacement (IWOR) sampling is an approximation-free technique for sharing permutation indices across a large number of permutation tests. An “IWOR sample” (top) is a single vector from which the set of permutation indices for each hypothesis can be extracted. The permutation indices for pair *A* consist of the first four elements of the IWOR sample; the permutation indices for pair *B* consist of the first five elements of the IWOR sample; and the permutation indices for pair *C* consist of the first six elements of the IWOR sample. Importantly, *the cells themselves are not shared across hypotheses*; rather, the IWOR sample is shared across hypotheses. IWOR sampling assumes that the number of control units is fixed across hypotheses; this assumption holds on low-MOI single-cell CRISPR screen data.

**Figure S11:**
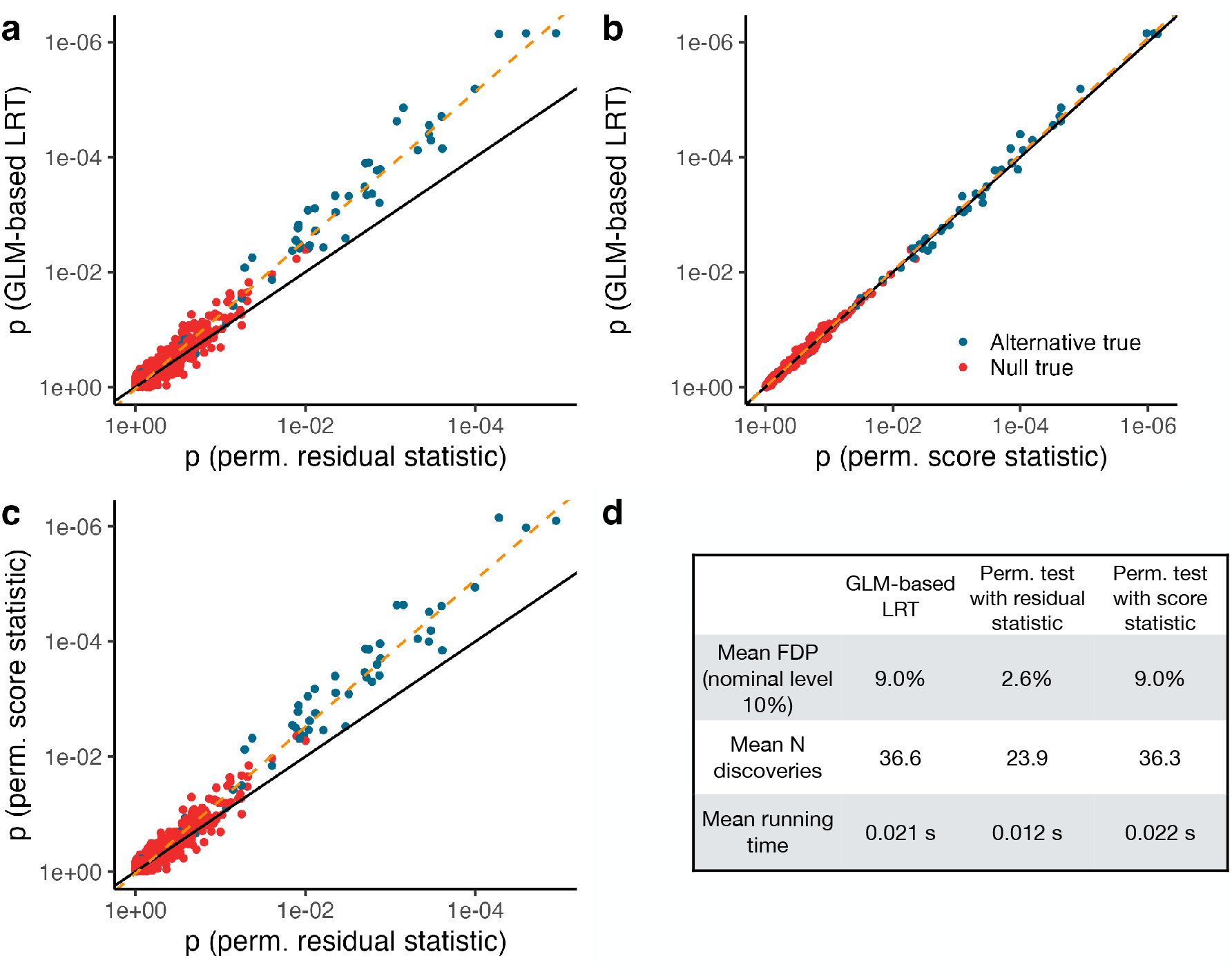
A comparison of the *T*_score_ and *T*_resid_ permutation test statistics on data simulated from an NB GLM. **a**, GLM likelihood ratio test (LRT) *p*-values vs. *T*_resid_-based permutation test *p*-values. The *T*_resid_-based *p*-values were generally larger (i.e., more conservative) than the GLM LRT *p*-values. **b**, GLM LRT *p*-values vs. *T*_score_-based permutation test *p*-values. These two sets of *p*-values were highly concordant. **c**, *T*_score_-based permutation test *p*-values vs. *T*_resid_-based permutation test *p*-values. The *T*_resid_-based permutation test *p*-values were generally more conservative than the *T*_score_-based permutation test *p*-values. **d**, mean false discovery proportion, number of discoveries, and running time of the GLM LRT, *T*_resid_-based permutation test, and *T*_score_-based permutation test across 500 replicates of the experiment. The GLM LRT and *T*_score_-based permutation test exhibited near-identical statistical and computational performance. The *T*_resid_-based permutation test exhibited lower power than the former two methods.

**Figure S12:**
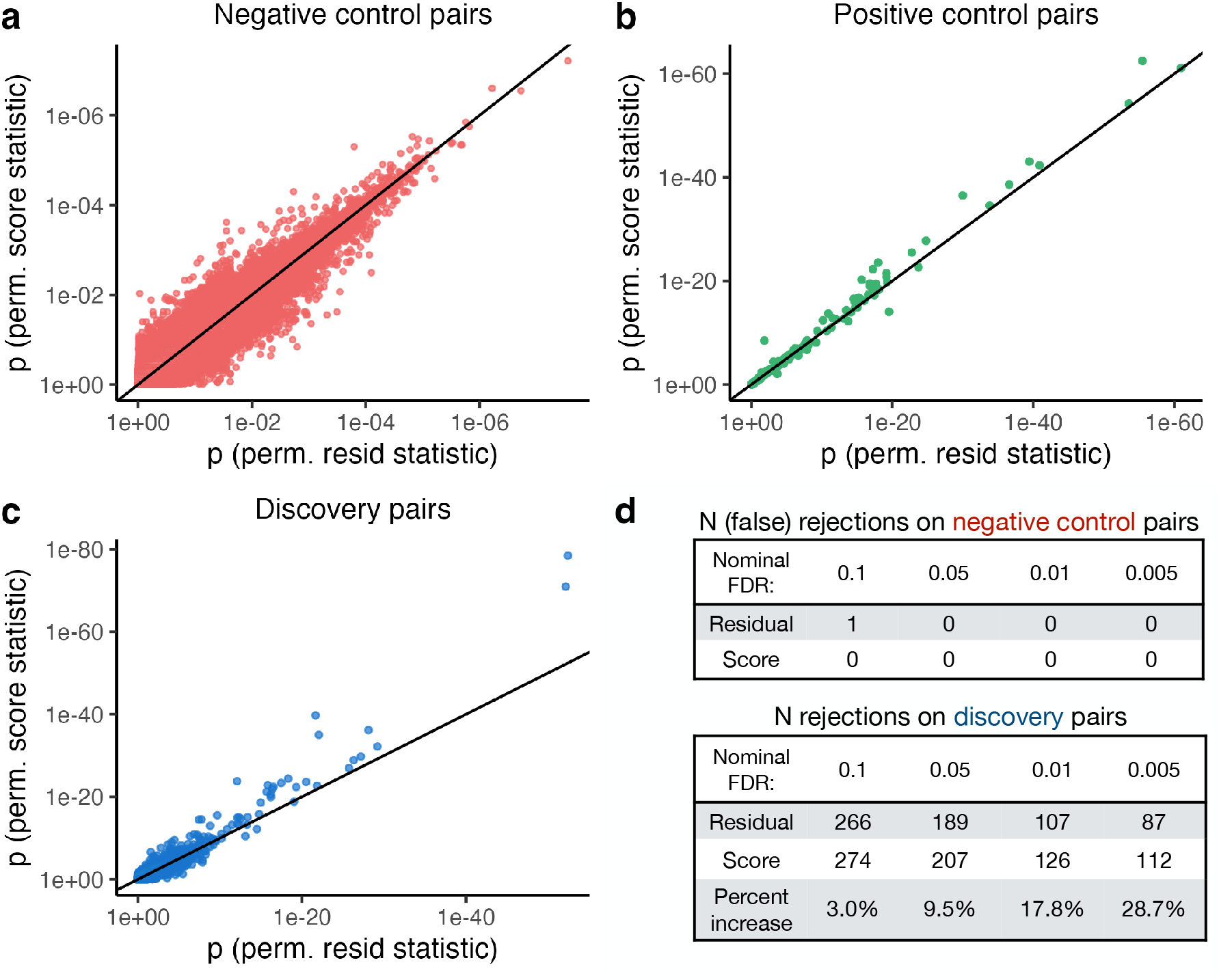
A comparison of the *T*_score_ and *T*_resid_ permutation test statistics on the Frangieh IFN-*γ* data. *T*_score_-based *p*-values vs. *T*_resid_-based *p*-values on the negative control (**a**), positive control (**b**), and discovery (**c**) pairs. **d**, number of rejections made by the *T*_resid_-based method (top row) and *T*_score_-based method (bottom row) on the negative control (top table) and discovery (bottom table) pairs. The various nominal FDR levels — 0.1, 0.05, 0.01, and 0.005 — are in the columns.

**Figure S13:**
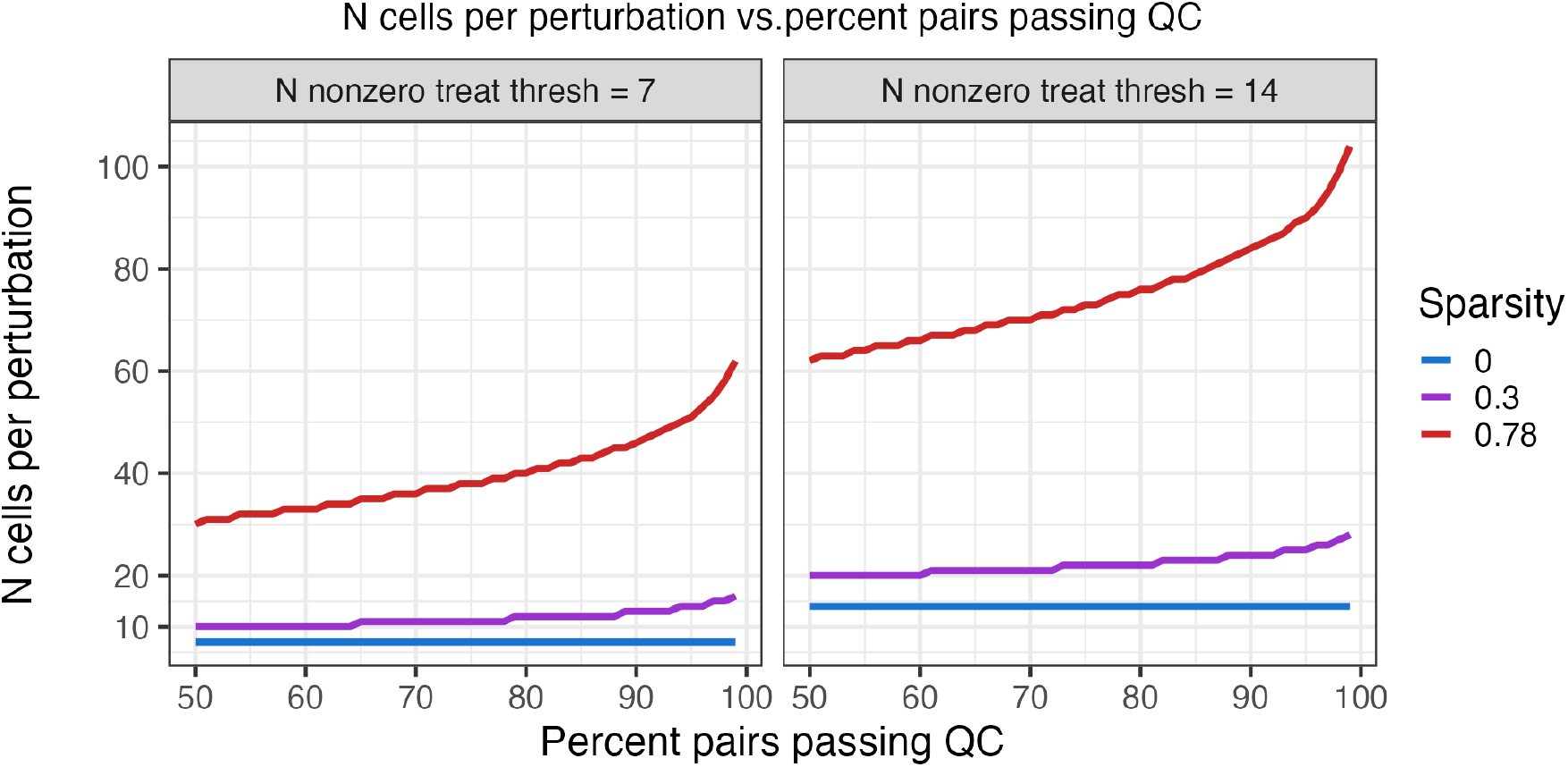
Number of cells per perturbation vs. probability that a given pair passes pairwise QC. A plot of the number of cells required per perturbation (*n*_trt_(*π, T, s*)) vs. the probability that a given pair passes pairwise QC (*π*). The left (resp., right) plot considers setting the pairwise QC threshold *T* to 7 (resp., 14). Finally, the colors indicate different sparsity levels *s*: red = 0.78, purple = 0.30, and blue = 0.0.

## S2 Additional mathematical details of SCEPTRE (low-MOI)

This section contains additional mathematical details of SCEPTRE (low-MOI). First, we derive the expression for the GLM score statistic used within the method, and next, we describe IWOR sampling.

### Derivation of the expression for the GLM score test statistic

We derive the GLM score test statistic for adding a single variable to a fitted GLM model. (We are not aware of any book or paper that presents this derivation.) Let *Y* ∈ ℝ^*n*^ be the vector of responses. Let ***Z*** ∈ ℝ^*n×p*^ be the matrix of variables onto which we are regressing *Y*. (Assume that ***Z*** includes a column of ones corresponding to the intercept.) Let *X* ∈ ℝ^*n*^ be the vector we seek to test for inclusion in the fitted GLM. Let ***D*** = [*X* ***Z***] ∈ ℝ^*n×*(*p*+1)^ be the augmented design matrix.

We model the mean *μ*_*i*_ of *Y*_*i*_ according to

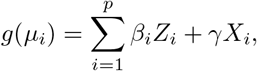

where *β* = [*β*_1_, …, *β*_*p*_]^*T*^ ∈ ℝ^*p*^ and *γ* are constants, and *g* : ℝ → ℝ is a link function. Let *θ* = [*β, γ*]^*T*^ ∈ ℝ^*p*+1^ denote the vector of unknown parameters. Also, let *η*_*i*_ = *g*^*−*1^(*μ*_*i*_) denote the linear component of the model, and let *V* (*μ*_*i*_) denote the variance of the response *y*_*i*_ given mean *μ*_*i*_. Standard GLM results indicate that the score *u* and Fisher information *I* of the model evaluated at *θ* are

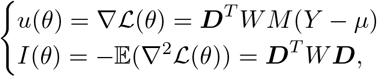

where *L*(*θ*) is the log-likelihood of the model evaluated at *θ*,

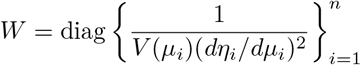

is the matrix of “weights” and

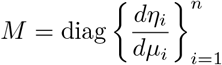

is the matrix of derivatives of the linear component with respect to the mean. We can rewrite the score as

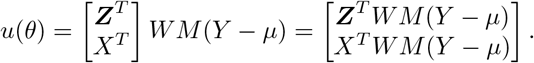

Next, we can rewrite the Fisher information as

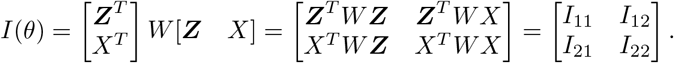

Define *I*_*X*_ by

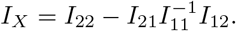

We have that

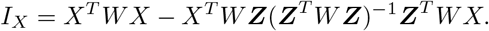

Let 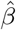 be the MLE for *β*. Next, let *u*_*X*_ denote the component of the score vector corresponding to *γ*, i.e.,

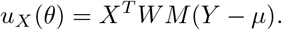

The value of *u*_*X*_ evaluated at 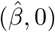 is

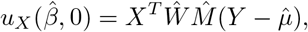

where *Ŵ*, 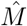, and 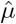 denote *W, M*, and *μ*, respectively, evaluated at 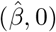. Similarly, the value of *I*_*X*_ (*θ*) evaluated at 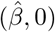 is

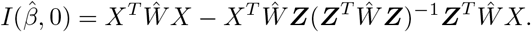

Standard asymptotic results indicate that

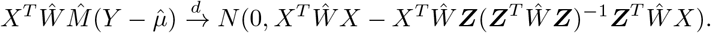

The GLM score test *z*-score is therefore

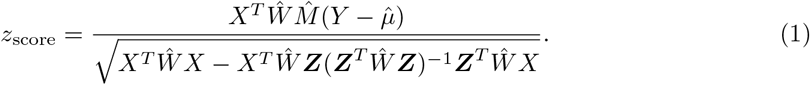

The GLM score test statistic typically is expressed in a different way in books and papers.[1] Let *E*_2_ denote the matrix of residuals after least squares regression of the columns of *X* onto ***Z***, i.e.,

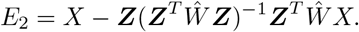

Furthermore, define 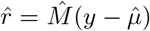 as the “working residual” vector. The GLM score test statistic typically is expressed as

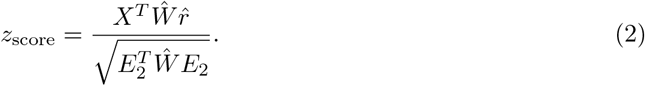

The expressions (1) and (2) are identical:

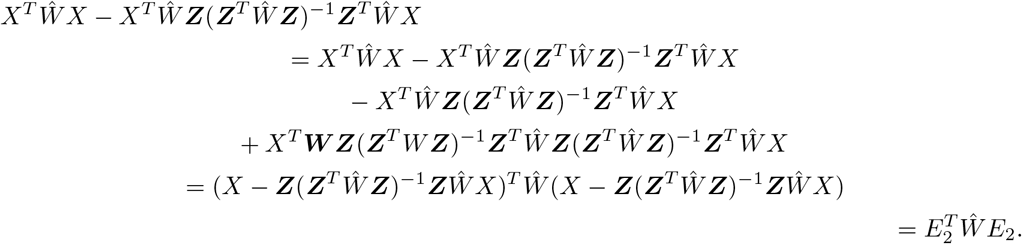

Expression (2) is evaluated via QR decomposition in practice. By contrast, we propose to evaluate (1) via spectral or Cholesky decomposition, thereby exploiting the sparsity of *X*.

### Inductive without replacement sampling

Permutation testing is a computationally expensive statistical technique. There are two main steps involved permutation testing: first, permuting the treatment vector *B* times, where *B* is some large integer (e.g., *B* = 5, 000), and second, recomputing the test statistic on the observed dataset and the *B* permuted datasets. Both of these steps are slow; which is slower depends on several factors, including the test statistic in use and the sparsity of the treatment vector. Permutation testing is even more expensive in the “high-throughput setting” in which one seeks to test tens of thousands (or more) of hypotheses via a permutation test. We introduce “inductive without replacement” (IWOR) sampling, a technique for recycling compute across permutation tests in which the number of “control units” is constant across tests (as in low-MOI single-cell CRISPR screens). IWOR sampling is a (provably) approximation-free technique, yielding an exact and correct permutation *p*-value for every hypothesis tested.

#### A concrete example of IWOR sampling

We first provide a concrete example of IWOR sampling to help convey intuition for the procedure. (We describe IWOR sampling in a mathematically precise way in the following section.) Suppose that we are analyzing a single-cell CRISPR screen dataset containing 19 cells, four of which contain a non-targeting perturbation, four of which contain targeting perturbation *A*, five of which contain targeting perturbation *B*, and six of which contain targeting perturbation *C*. Suppose that we seek to test for association between each of the targeting perturbations and some gene. Call pair *A* (resp., pair *B*, pair *C*) the pair that results from coupling perturbation *A* (resp., perturbation *B*, perturbation *C*) to the gene.

Figure S10 depicts the “observed treatment vector” corresponding to pair *A*, pair *B*, and pair *C*. For example, the observed treatment vector for pair *A* contains four treatment cells (blue circles) and four control cells (gray circles), while the observed treatment vector for pair *B* contains five treatment cells and four control cells, etc. We can enumerate the cells contained within each pair by assigning an integer index to each cell. For example, we can enumerate the cells contained within pair *A* by 1, …, 8, and we can enumerate the cells contained within pair *B* by 1, …, 9, etc. (Note: although we reuse integer indices across pairs, the cells themselves *are not shared* across pairs.) We seek to permute the observed treatment vector corresponding to each pair *B* times. This is equivalent to drawing *B* without replacement samples of size *k* from the set of indices 1, …, *k* + *N*, where *k* is the number of treatment cells in a given pair and *N* is the number of control cells in the dataset. For example, consider the observed treatment vector corresponding to pair *A* (Figure S10, left column). We can permute this vector by assigning cells 2, 4, 5, and 6 to “treatment” status and assigning the remaining cells to “control” status. Moreover, we can identify this particular permutation of the vector with the set of integers {2, 4, 5, 6}, where {2, 4, 5, 6} can be thought of as a without-replacement sample from the set of indices 1, …, 8. Here, 8 is the sum of the number of control cells (4) and the number of treatment cells (4).

Our goal is to randomly permute the observed treatment vector corresponding to each pair. IWOR sampling is a strategy for doing this efficiently. An example IWOR sample for this dataset is *v* := [5, 6, 4, 2, 9, 10] (Figure S10, top). The set that contains the first *k* elements of *v* constitutes a valid without-replacement sample of size *k* from the set of indices 1, …, *k* + 4. (The significance of 4 in this context is that 4 is the number of control cells in the example dataset.) For example, the set constructed from the first four elements of *v* — {5, 6, 4, 2} — is a without-replacement sample from the set 1, …, 8. Thus, {5, 6, 4, 2} is a valid permutation of the treatment vector corresponding to pair *A*, which contains four treatment cells.

Next, the set containing the first five elements of *v* — {5, 6, 4, 2, 9} — is a valid without-replacement sample from the set 1, …, 9. Thus {5, 6, 4, 2, 9} is a valid permutation of the treatment vector corresponding to pair *B*, which contains five treatment cells. Finally, the set constructed from the first six elements of *v* — *v* itself — is a valid without replacement sample from the set 1, …, 10. Thus, *v* is a valid permutation of the treatment vector corresponding to pair *C*, which contains six treatment cells. To permute each treatment vector *B* times, we generate *B* IWOR samples. For example, four possible IWOR samples for the above dataset are as follows: [5, 6, 4, 2, 9, 10], [4, 5, 7, 8, 2, 9], [5, 2, 1, 8, 3, 10], and [3, 4, 7, 1, 2, 10].

More generally, suppose a given dataset contains *N* control cells. Let *x* be a random (i.e., unrealized) IWOR sample, and let *x*[1 : *k*] denote the first *k* elements of *x*. The I WOR s ample is carefully constructed such that it satisfies the following crucial property:

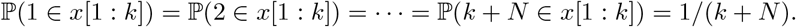

In other words, the first *k* elements of *x* constitute a valid WOR sample from the set {1, …, *k* +*N*}. The permutation defined by *x*[1 : *k*] is therefore a valid permutation for a pair containing *k* treatment cells and *N* control cells.

#### Definition o f I WOR sampling

We provide a more precise definition of IWOR sampling here. Consider a gene expression vector *Y* ∈ ℝ^*n*^, technical factor matrix ***Z*** ∈ ℝ^*n×p*^, and binary “treatment” vector of perturbation indicators *X* ∈ {0, 1}^*n*^. We can identify a given treatment vector with the set of nonzero entries within that vector. In other words, we can represent the vector *X* as the set *S* = {*i* : *X*[*i*] = 1}, where *X*[*i*] denotes the value of the vector *X* at position *i*. Suppose that *X* has *m* nonzero entries. Randomly permuting *X* is equivalent to sampling *m* elements without replacement (WOR) from the set {1, …, *n*}. Unfortunately, drawing *B* WOR samples for every perturbation-gene pair — which is required for carrying out the permutation test dataset-wide — is slow. IWOR sampling shares WOR samples across pairs.

A key observation is that the number of control cells is fixed across all hypotheses, while the number of treatment cells — i.e., the number of cells containing the targeting perturbation — varies from hypothesis to hypothesis (depending on the targeting perturbation). We exploit this structure of the problem as follows. Let *N* be the number of control cells; label the control cells by *c*_1_, *c*_2_, …, *c*_*N*_. Assume that there are *p* targeting perturbations. For perturbation *i* ∈ {1, …, *p*}, let *k*_*i*_ be the number of cells containing perturbation *i*. Finally, let *M* = max_*i*∈{1,…,*p*}_ *k*_*i*_ be the number of cells containing the perturbation that infects the greatest number of cells. Label the treatment cells by *t*_1_, …, *t*_*M*_. We construct a length-*M* random sequence *a*_1_, *a*_2_, …, *a*_*M*_ that satisfies the following three properties:

1. *a*_*i*_ ∈ {*c*_1_, …, *c*_*N*_, *t*_1_, …, *t*_*i*_} for all *i* ∈ {1, …, *M*}; that is, the *i*th element of the sequence is a control cell or one of the first *i* treatment cells.
2. *a*_*i≠*_ *a*_*j*_ for *i* ≠ *j*. That is, the elements of the sequence are unique. Let *A*_*i*_ := {*a*_1_, …, *a*_*i*_} denote the set containing the first *i* elements of the sequence. (Observe that there are *N* + *i* elements that possibly could be in *A*_*i*_: the *N* control cells and the first *i* treatment cells.) We additionally require the following property to hold:
3. ℙ (*c*_1_ ∈ *A*_*i*_) = *· · ·* = ℙ (*c*_*N*_ ∈ *A*_*i*_) = ℙ (*t*_1_ ∈ *A*_*i*_) = *· · ·* = ℙ (*t*_*i*_ ∈ *A*_*i*_) = *i/*(*i* + *N*). That is, each of the elements that possibly could be in *A*_*i*_ has an equal chance of being in *A*_*i*_.

The sequence *a*_1_, …, *a*_*M*_ yields a sequence of increasing sets *A*_1_ ⊂ *A*_2_ ⊂ *· · ·* ⊂ *A*_*m*_. The first set *A*_1_, which contains a single element, contains the control cells and the first treatment cell with equal probability. The second set *A*_2_, which contains two elements, contains the control cells and each of the first *two* treatment cells with equal probability, and so on. Thus, if a given gene-perturbation pair contains *k* ≤ *M* treatment cells, then *A*_*k*_ constitutes a valid WOR sample for that pair. Importantly, because the sequence of sets *A*_1_, …, *A*_*M*_ is increasing, we can share the random sample *a*_1_, …, *a*_*M*_ across all gene-perturbation pairs. We call this technique “inductive without replacement sampling” (or “IWOR sampling”). The SCEPTRE (low MOI) algorithm generates a set of *B* length-*M* IWOR samples and shares this set of samples across all pairs.

#### An abstract approach for constructing an IWOR sample

We describe an abstract approach for constructing an IWOR sample and prove its correctness. The procedure is inductive.

**Step 1**. Sample one element from the set {*c*_1_, …, *c*_*N*_, *t*_1_}, putting an equal mass of 1*/*(*N* + 1) onto each of the elements. Call the sampled element *a*_1_. Set *A*_1_ = {*a*_1_}.

**Step** *i* **for** *i* ∈ {2, …, *M*}. Let *B*_*i*_ := {*c*_1_, …, *c*_*N*_, *t*_1_ …, *t*_*i−*1_} *\ A*_*i−*1_ denote the set of “leftover” elements not sampled in steps 1, …, *i −* 1. There are *N* elements in the set *B*_*i*_. Draw an element at random from the set *B*_*i*_ ⋃ {*t*_*i*_}, placing a mass of 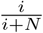 on *t*_*i*_ and a mass of 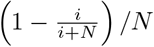 on each of the elements in *B*_*i*_. Call the sampled element *a*_*i*_. Set *A*_*i*_ = *A*_*i−*1_ ∪ {*a*_*i*_}.

The resulting sequence *a*_1_, …, *a*_*M*_ satisfies the three properties listed above, which we now prove.

**Proof** : It is clear that *a*_*i*_ ∈ {*c*_1_, …, *c*_*N*_, *t*_1_, …, *t*_*i*_} for all *i* and that *a*_*i*_ ≠ *a*_*j*_ for *i* ≠ *j*. We focus on proving the third property, which we do inductively.

Base case: Let *i* = 1. Then

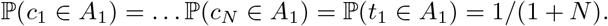

Inductive step: Suppose that

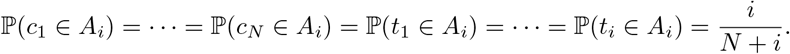

To construct the set *A*_*i*+1_, we sample *t*_*i*+1_ with probability (*i* + 1)*/*(*N* + *i* + 1) and each element of *B*_*i*_ with probability

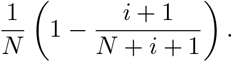

We call the sampled element *a*_*i*+1_, and we set *A*_*i*+1_ = *A*_*i*_ ∪ {*a*_*i*+1_}. Our goal is to show that, for all *u* ∈ {*c*_1_, …, *c*_*N*_, *t*_1_, …, *t*_*i*+1_},

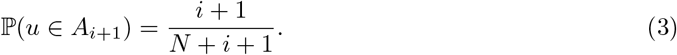

(That is, for all elements that *could* be in *A*_*i*+1_, each has an equal chance of *actually being* in *A*_*i*+1_.) The equality (3) holds for *u* = *t*_*i*+1_ by construction. Next, suppose that *u* ∈ {*c*_1_, …, *c*_*N*_, *t*_1_, …, *t*_*i*_}. We compute ℙ (*u* ∈ *A*_*i*+1_) using the law of total probability:

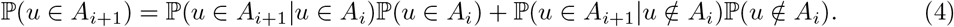

We consider first the term ℙ (*u* ∈ *A*_*i*+1_|*u* ∈ *A*_*i*_)P(*u* ∈ *A*_*i*_). We have that ℙ (*u* ∈ *A*_*i*+1_|*u* ∈ *A*_*i*_) = 1, as membership of *u* in *A*_*i*_ implies membership of *u* in *A*_*i*+1_. By the inductive hypothesis, ℙ (*u* ∈ *A*_*i*_) = *i/*(*N* + *i*), implying that

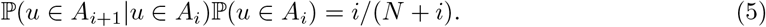

Next, we consider the term ℙ (*u* ∈ *A*_*i*_+1|*u* ∉ *A*_*i*_) ℙ (*u* ∉ *A*_*i*_). If *u* ∉ *A*_*i*_ (i.e., if *u* is a “leftover” element at step *i*), then *u* ∈ *B*_*i*_, implying

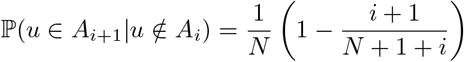

by construction. Furthermore, by the inductive hypothesis, ℙ (i ∉ Ai) = 1−ℙ (*u* ∈ *A*_*i*_) = 1−*i*/(*N* +*i*).. Thus,

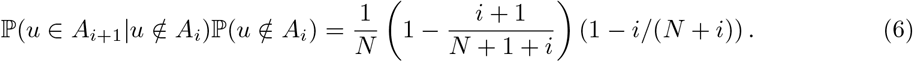

Combining (4), (5), and (6),

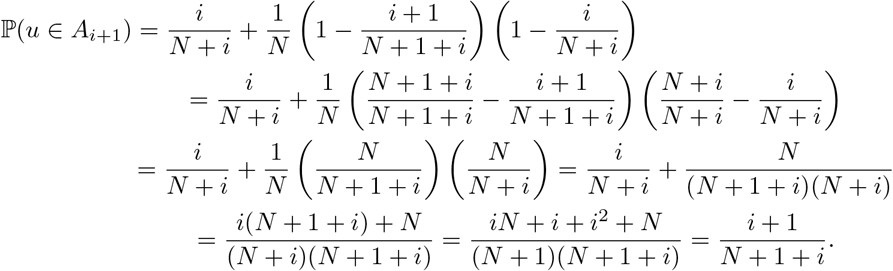

#### A concrete algorithm for IWOR sampling

We now derive a concrete algorithm for constructing an IWOR sample of size *M* on the set {*c*_1_, …, *c*_*N*_, *t*_1_, …, *t*_*M*_}. The algorithm requires only a high-quality uniform random number generator, which is readily available in most programming languages. First, we describe a simple algorithm for sampling from the discrete probability distribution with support {0, …, *N*} that places mass *i/*(*i* + *N*) on *N* and mass (1*/N*)[1 *− i/*(*i* + *N*)] on {0, 1, …, *N −* 1}. (We call this distribution the `IWOR`(*N, i*) distribution.) The algorithm, given in Algorithm 1, takes *O*(1) time and space.

Next, we provide an algorithm for constructing an IWOR sample of length *M* on the set of control cells {*c*_1_, …, *c*_*N*_} and treatment cells {*t*_1_, …, *t*_*M*_}. Algorithm 2 is a concrete instantiation of the “abstract approach for constructing an IWOR sample,” as described in the previous section. The vector *r* is a vector of length *N* + 1. At step *i*, the *N* cells that remain (i.e., those that have not yet been sampled among {*c*_1_, …, *c*_*N*_, *t*_1_, …, *t*_*i−*1_}) are stored in the leftmost *N* positions of *r*. The cell *t*_*i*_ is then inserted into the rightmost position of *r*. A cell is sampled from *r* via the `IWOR`(*N, i*) distribution, which places more mass on the cell in the rightmost position (i.e., *t*_*i*_). The sampled cell is moved into *v*. Then, the cell in the rightmost entry of *r* (i.e., *t*_*i*_) is moved into the position vacated by the sampled cell. Finally, the cell *t*_*i*+1_ is moved into the rightmost position of *r*. We

##### Algorithm 1

Generating a sample from the `IWOR`(*N, i*) distribution.

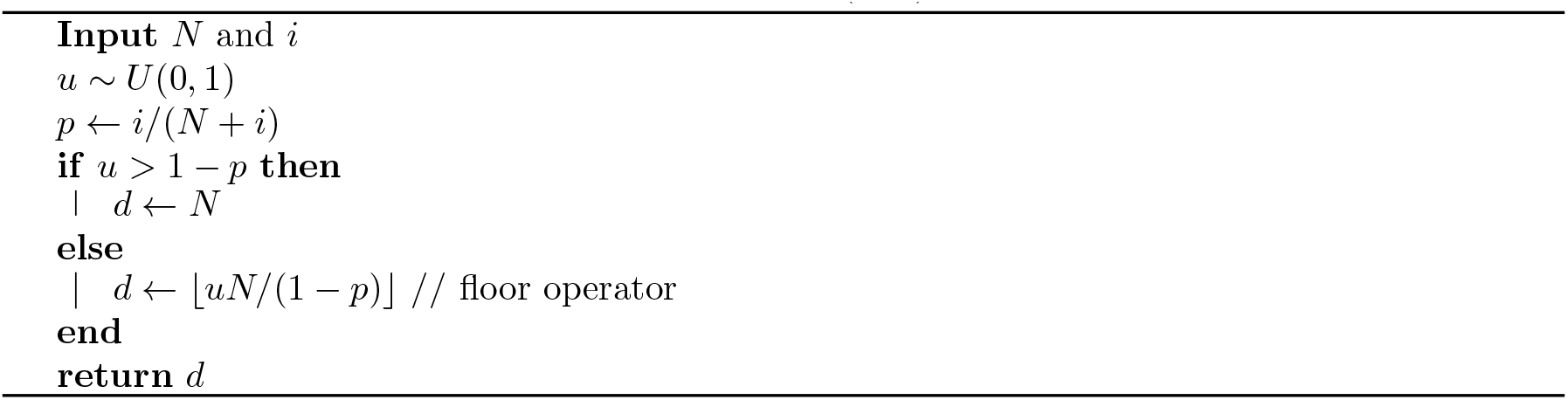

repeat this process until we have iterated through all of the treatment cells. This algorithm is fast and efficient, taking *O*(*M*) time and *O*(*M* + *N*) space.

##### Algorithm 2

Generating an IWOR sample of size *M* given control cells *c*_1_, …, *c*_*N*_ and treatment cells *t*_1_, …, *t*_*M*_.

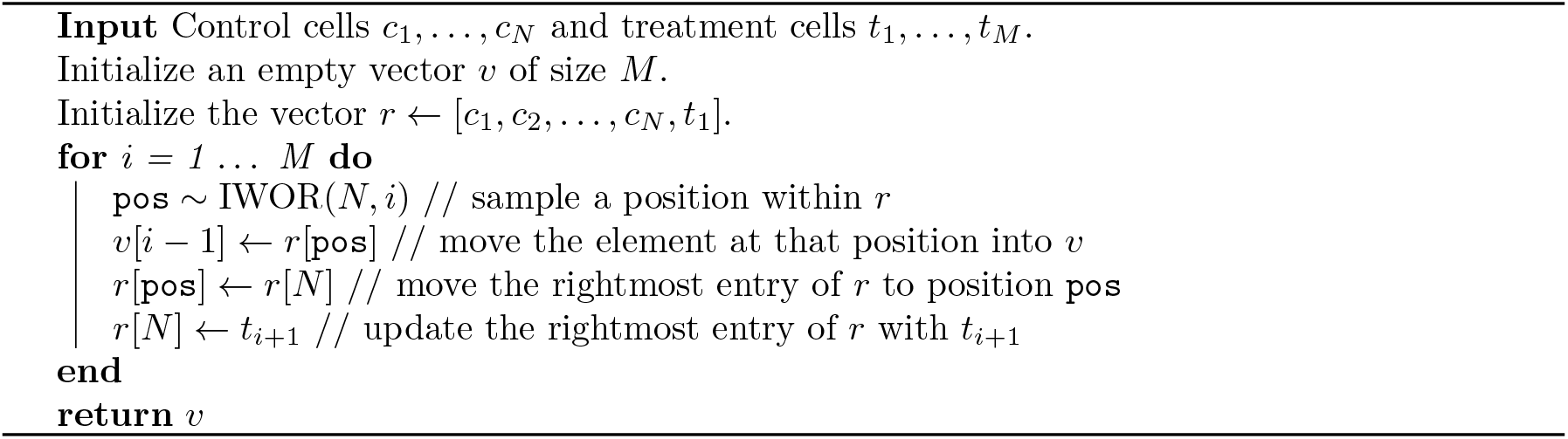

SCEPTRE (low-MOI) uses a slightly modified, more efficient, and more numerically stable version of Algorithm 2 for constructing the IWOR sample. Suppose there are *p* targeting perturbations, with the *i*th perturbation infecting *k*_*i*_ cells. Let *M* = max_*i*∈{1,…,*p*}_ *k*_*i*_ be the number of cells containing the perturbation that infects the greatest number of cells, and let *m* = min_*i*∈{1,…,*p*}_ *k*_*i*_ be the number of cells containing the perturbation that infects the least number of cells. SCEPTRE first samples *k* cells without replacement from the set *c*_1_, …, *c*_*N*_, *t*_1_, …, *t*_*k*_. Then, SCEPTRE constructs an IWOR sample, taking as input the cells that remain from the first step and treatment cells *t*_*k*+1_, *t*_*k*+2_, …, *t*_*M*_.

## S3 Additional empirical analyses

In this section we present several additional empirical analyses.

### Comparing the score statistic to the difference-in-residual-means statistic

SCEPTRE (low-MOI) uses a permutation test based on an NB GLM score statistic. A natural question is whether SCEPTRE (low-MOI) is better (in some sense) than the following alternative method: first, “regress out” the covariates by regressing the response vector onto the covariate matrix via an NB GLM; second, extract the residuals^1^ from the fitted GLM; third, perform a permutation test between the treatment vector (i.e., the vector of perturbation presences/absences) and the vector of residuals, taking as the test statistic the standard difference in means statistic. Using mathematical language, let *Y* = [*Y*_1_, …, *Y*_*n*_]^*T*^ be the vector of gene (or protein, etc.) UMI counts, *X* = [*X*_1_, …, *X*_*n*_]^*T*^ the vector of perturbation presence or absence indicators, and *Z* ∈ R^*n×p*^ the matrix of technical factors (or covariates). Regressing *Y* onto *Z* via an NB GLM yields a vector of fitted means 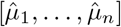. Define the *i*th (raw) residual by 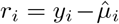. The difference-in-residual-means test statistic *T*_resid_ is defined as

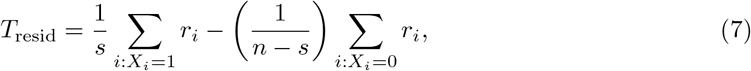

where 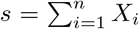 is the number of “treated” cells (i.e., the number of cells containing a perturbation). The GLM score statistic, meanwhile, is defined as

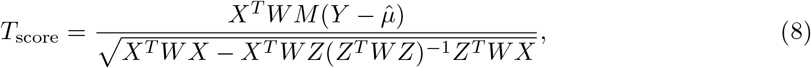

where *W* and 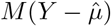 are given by

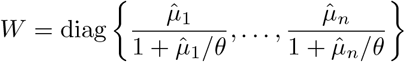

and

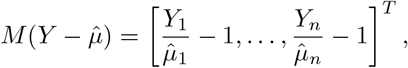

respectively. Clearly, *T*_score_ is more complex than *T*_resid_: evaluating the denominator of the former involves performing matrix multiplication, addition, transposition, and inversion operations. How (if at all) do *T*_score_ and *T*_resid_ differ with respect to the *p*-values that they produce? To explore this question, we applied a permutation test based on *T*_score_ and another based on *T*_resid_ to real and simulated data, comparing the two methods with respect to type-I error control, power, and computational performance. We found that *T*_score_ and *T*_resid_ exhibited similar type-I error control and computational performance but that *T*_score_ could be much more powerful than *T*_resid_. We note that these results — while interesting — are preliminary; we intend to carry out a more in-depth comparison between *T*_score_ and *T*_resid_ in a followup, more statistically-focused work.

#### Simulation study

We first compared *T*_resid_ to *T*_score_ on simulated data. We used *n* = 5, 000 cells in the simulation study. We constructed the cell-specific covariates *z*_1_, …, *z*_*n*_ ∈ ℝ^3^ by drawing *n* i.i.d. samples from a bivariate normal distribution and appending a one for the intercept term. (The cell-specific covariates were meant to represent technical factors, such as sequencing depth and percent mitochondrial reads.) Next, we generated the *i*th binary treatment indicator *x*_*i*_ by sampling *x*_*i*_ from a Bernoulli distribution with mean *μ*_*i*_ = *σ*^*−*1^(*τ* ^*T*^ *z*_*i*_), where *σ* was the logit function and *τ* ∈ ℝ^3^ a fixed constant. Thus, the treatment *x*_*i*_ and covariate *z*_*i*_ were correlated, putting us into the *confounded* (as opposed to *unconfounded*) problem setting.

We then simulated gene expression data on *n*_genes_ = 500 genes. For a given gene *j* ∈ {1, …, *n*_genes_}, we sampled UMI counts 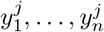 from an NB GLM model:

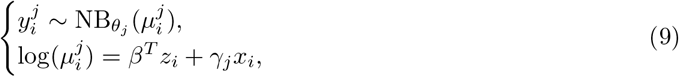

where *β*^*T*^ ∈ ℝ^*p*^ was a fixed regression coefficient, *θ*_*j*_ *>* 0 was a gene-specific NB size parameter, and *γ*_*j*_ ∈ ℝ was a gene-specific treatment effect. To simulate variation in overdispersion across genes, we sampled 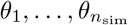 from a Unif(0.1, 5) distribution. Additionally, to simulate variation in response to the treatment, we set the gene-specific treatment effect *γ*_*j*_ to 0 for 450 (randomly-selected) genes, and we set *γ*_*j*_ to a small, positive constant for the remaining 50 genes. Thus, for 450 (or 90% of) genes, we simulated data from the *null* model (in which treatment and response *were not* associated), and for 50 (or 10% of) genes, we simulated data from an *alternative* model (in which treatment and response *were* associated).

We applied three methods to analyze the data: a permutation test based on the difference-in-residual-means statistic *T*_resid_, a permutation test based on the score test statistic *T*_score_, and a GLM likelihood ratio test. We employed the adaptive permutation testing scheme and skew-normal approximation of SCEPTRE (low-MOI) to accelerate the two permutation-based methods (see Acceleration 3: Adaptive permutation testing and Acceleration 4: Skew-normal fit in the Methods section for details). Additionally, to simplify comparison of the methods, we treated the NB size parameter *θ*_*j*_ as fixed and known. (We estimated the NB size parameter in the real data analysis; see next section.) The GLM likelihood ratio test consisted of comparing the full model (9) to the reduced model that results from setting *γ*_*j*_ to zero in (9). We note that the GLM likelihood ratio test was optimal in the simulation study setting (i.e., the setting in which the GLM is correctly specified and the sample size is sufficiently large so as to enable accurate asymptotic inference). The *T*_resid_-based permutation test and *T*_score_-based permutation test were applied with two-sided test statistics so as to enable a meaningful comparison with the GLM likelihood ratio test, which is an inherently two-tailed test.

We assessed the degree of concordance between the *T*_score_-based permutation test *p*-values, the *T*_resid_-based permutation test *p*-values, and the GLM likelihood ratio test (LRT) *p*-values. As the GLM LRT *p*-values were optimal, we looked for agreement between the GLM LRT *p*-values and the *p*-values outputted by the two permutation methods. First, we plotted the GLM LRT *p*-values against the *T*_resid_-based permutation test *p*-values (Figure S11a). Each point in the plot represents one of the 500 hypotheses tested; the red (resp., blue) points correspond to the null (resp., alternative) hypotheses. The vertical (resp., horizontal) position of a given point indicates the GLM LRT (resp., *T*_resid_-based permutation test) *p*-value computed on that hypothesis. The *p*-values were negative log-transformed to facilitate visualization of small *p*-values. We observed that the *T*_resid_-based *p*-values were strongly and positively correlated with the GLM LRT *p*-values. However, the *T*_resid_-based *p*-values were generally larger (i.e., less significant) than the GLM LRT *p*-values. We regressed the GLM LRT *p*-values onto the *T*_resid_-based *p*-values (via OLS) and superimposed the line of best fit onto the plot (dashed orange line). The line of best fit clearly lay above the identity line (solid black line), indicating deflation of the *T*_resid_-based *p*-values relative to the GLM LRT *p*-values.

Next, we plotted the GLM LRT *p*-values against the *T*_score_-based permutation *p*-values (Figure S11b). The two sets of *p*-values were highly concordant, as evidenced by the tight clustering of the points around the identity line (solid black line). Furthermore, the line of best fit (solid orange line) closely tracked the identity line, indicating a high degree of agreement between the two sets of *p*-values. Finally, we plotted the *T*_score_-based permutation *p*-values against the *T*_resid_-based permutation *p*-values (Figure S11c). As expected, the *T*_score_-based *p*-values were generally smaller (i.e., more significant) than the *T*_resid_-based *p*-values. We concluded that the *T*_score_-based permutation *p*-values demonstrated a higher degree of concordance with the GLM LRT *p*-values than did the *T*_resid_-based permutation *p*-values.

Finally, we subjected the three sets of *p*-values (i.e., those produced by the *T*_score_-based permutation test, the *T*_resid_-based permutation test, and the GLM LRT) to a BH correction at level 10%, obtaining a discovery set for method. The GLM LRT yielded 41 discoveries, two of which were false discoveries (false discovery proportion = 4.9%); next, the *T*_score_-based permutation test produced 42 discoveries, two of which were false discoveries (false discovery proportion = 4.7%); finally, the *T*_resid_-based permutation test yielded 25 discoveries and zero false discoveries (false discovery proportion = 0%). All three methods controlled the false discovery proportion at the nominal level of 10%. However, the *T*_score_-based permutation test and GLM LRT yielded about 68% more discoveries than the *T*_resid_-based permutation test.

To ensure robustness of the above results to choice of the random seed, we repeated the above experiment 500 times (Figure S11d). Averaged across runs, the GLM LRT, the *T*_score_-based permutation test, and the *T*_resid_-based permutation test yielded a mean false discovery proportion of 9.0%, 9.0%, and 2.6%, respectively. (The mean false discovery proportion functioned as an estimate for the false discovery rate.) Thus, all methods controlled the false discovery rate at the nominal level of 10%.^2^ Next, the GLM LRT, the *T*_score_-based permutation test, and the *T*_resid_-based permutation test yielded a mean of 36.6 discoveries, 36.3 discoveries, and 23.9 discoveries, respectively. Hence, the GLM LRT and *T*_score_-based permutation test produced about 53% more discoveries than the *T*_resid_-based permutation test. Finally, the GLM LRT, the *T*_score_-based permutation test, and the *T*_resid_-based permutation test had a mean per-hypothesis compute time of 0.021 s, 0.022 s, and 0.012 s, respectively. Thus, the GLM LRT and *T*_score_-based permutation test demonstrated similar running times, while the *T*_resid_-based permutation test was roughly twice as fast.

In summary the GLM LRT and the *T*_score_-based permutation test exhibited nearly identical performance in the simulation study: the two methods controlled type-I error at the same rate, made virtually the same number of discoveries, and had essentially the same running time. The *T*_resid_-based permutation test, on the other hand, was considerably less powerful than either the GLM LRT or the *T*_score_-based permutation test. We highlight that — in principle — the *T*_score_-based permutation test can retain validity in a much broader range of settings than the standard GLM LRT, such as settings in which the data are sparse or the NB model is misspecified. We conjecture that the *T*_score_-based permutation test trades a small loss in power for a substantial improvement in robustness relative to a standard GLM LRT. Additional work remains to be done to understand this phenomenon.

#### Real data analysis

We applied both the *T*_score_-based permutation test and the *T*_resid_-based permutation test to the Frangieh IFN-*γ* data. Similar to the simulation study, both methods controlled type-I error, while the *T*_score_-based test produced a larger discovery set than the *T*_resid_-based test. We analyzed three sets of perturbation-gene pairs: negative control pairs, positive control pairs, and discovery pairs. The discovery pairs consisted of the entire set (i.e., the *trans* set) of perturbation-gene pairs (that passed pairwise QC). The negative control pairs were constructed by the SCEPTRE software and were “matched” to the discovery pairs.^3^ In particular, each negative control target consisted of three, randomly-grouped non-targeting gRNAs. (In this sense the calibration check analysis reported here is not precisely the same as the calibration check analysis reported in Figure 4.) Finally, the positive control pairs consisted of perturbations targeting gene transcription start sites paired to the genes regulated by those transcription start sites.

Initially, the *p*-values outputted by the *T*_score_-based method and the *T*_resid_-based method exhibited mild miscalibration on the negative control data: the former method yielded five false discoveries, and the latter method yielded three false discoveries (at BH level 0.1). Therefore, we slightly increased the stringency of the cellwise QC thresholds, removing cells whose percent mitochondrial value exceeded 0.1 or whose number of genes expressed or total gene UMI count fell outside of the [0.1, 0.99] percentile range. We plotted the resulting *T*_score_-based *p*-values against the *T*_resid_-based *p*-values computed on the negative control pairs (S12a), positive control pairs (S12b), and discovery pairs (S12c). We subjected both the *T*_score_-based *p*-values and the *T*_resid_-based *p*-values to a BH correction (separating out the negative control pairs, positive control pairs, and discovery pairs). We varied the nominal FDR level over the grid {0.1, 0.05, 0.01, 0.005} to explore how increasing the stringency of the multiplicity correction might affect the results. The *T*_score_-based negative control *p*-values and *T*_resid_-based negative control *p*-values were roughly symmetrically distributed about the identity line (S12a). Moreover, both the *T*_score_-based test and *T*_resid_-based test controlled type-I error on the negative control pairs, as each method produced zero or one false discoveries at the various BH levels (S12d).

Next, we observed that the positive control *T*_score_-based *p*-values and *T*_resid_-based *p*-values were highly correlated (S12b). However, the *T*_score_-based *p*-values were generally smaller (i.e., more significant than) their *T*_resid_-based counterparts, hinting at the greater power of the *T*_score_ test statistic. Finally, the discovery *T*_score_-based *p*-values and *T*_resid_-based *p*-values also were highly correlated, and again, the *T*_score_-based *p*-values were generally smaller than their *T*_resid_ counterparts, especially in the tail of the *p*-value distribution. Moreover, the *T*_score_-based method yielded a larger discovery set than the *T*_resid_-based method over the various nominal FDR levels. The relative difference between the size of the of the *T*_score_-based discovery set and *T*_resid_-based discovery set increased as the BH correction became more stringent. For example, at BH level 0.1, the *T*_score_-based method yielded 274 discoveries, a 3% improvement over the *T*_resid_-based method, which made 266 discoveries. However, at BH level 0.005, the *T*_score_-based method produced 112 discoveries, a 29% improvement over the *T*_resid_-baesd method, which made 87 discoveries (Figure S12d).

We repeated this analysis on the Papalexi gene expression data, obtaining broadly similar results: the *T*_score_-based and *T*_resid_-based methods both controlled type-I error on the negative control data, but the *T*_score_-based method produced slightly smaller *p*-values on the positive control pairs, and the *T*_score_-based method yielded a larger discovery set (by about 3%) on the discovery pairs. Overall, synthesizing the results of the simulation study, the Frangieh data analysis, and the Papalexi data analysis, we concluded that the *T*_score_ statistic and *T*_resid_ statistic can yield quite different results. In particular, depending on the dataset under analysis and the stringency of the multiple testing adjustment, the *T*_score_ statistic can produce a discovery set up to 50% larger than the one produced by the *T*_resid_ statistic.

### Comparing the spectral decomposition algorithm to the QR decomposition algorithm for computing GLM score tests

We proposed a spectral decomposition algorithm for computing GLM score tests within the context of SCEPTRE (see Acceleration 2: A fast score test for binary treatments). The spectral decomposition approach differs from the standard approach for computing GLM score tests; the latter is based on a QR decomposition. We conducted a small computational experiment to compare the speed of the spectral decomposition algorithm to that of the QR decomposition algorithm. We found that the former was faster for sparse, binary treatment vectors. Furthermore, as we increased the sparsity level of the treatment vector, we found that the speed of the spectral decomposition algorithm relative to that of the QR decomposition algorithm increased.

We generated synthetic data on *n* = 100, 000 cells, drew the covariates *z*_1_, …, *z*_*n*_ from a bivariate Gaussian distribution, and generated the gene expression counts from a negative binomial model:

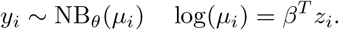

We regressed the gene expression counts onto the covariates, obtaining a fitted GLM fit.

Next, for a given sparsity level *π* ∈ {0.999, 0.99, 0.95, 0.90, 0.50}, we generated 1000 synthetic treatment vectors *x*^1^, …, *x*^1000^ (each of length *n*) by sampling i.i.d. variates from a Bern(1 *− π*) distribution. We tested each treatment vector for inclusion in the fitted GLM `fit` by running a GLM score test. We evaluated two different methods for computing the GLM score test: the spectral decomposition approach (proposed in this work), and the standard QR decomposition approach. We used the implementation of the QR decomposition approach available within the `statmod` [1] R package, which is a fast and high-quality implementation of the method. To ensure robustness of the results to choice of the random seed, we repeated the experiment 50 times for each choice of the sparsity level *π*, generating fresh covariates *z*_1_, …, *z*_*n*_ and gene expression counts *y*_1_, …, *y*_*n*_ for each run of the experiment.

We found that the spectral decomposition algorithm was much faster than the QR decomposition algorithm (Table S4). For example, at sparsity level 99.9%, the mean running time of the spectral decomposition algorithm was 0.0057*s* (95% CI: (0.0056*s*, 0.0058*s*)), while that of the QR decomposition algorithm was 1.55*s* (95% CI: (1.55*s*, 1.56*s*)), indicating a speedup by a factor of 272. Consistent with theory, the relative efficiency of the spectral decomposition algorithm in comparison to the QR decomposition algorithm decreased as the sparsity level decreased. For example, at sparsity level 50%, the spectral decomposition algorithm was only 11 times faster than the QR decomposition algorithm. We concluded that the spectral decomposition algorithm is well-suited to computing GLM score tests for sparse, binary treatment vectors.

### Experimental design considerations

A question that an experimenter might ask is as follows: “How many cells per perturbation do I need if I plan to use SCEPTRE to analyze my data?” We sought to provide a preliminary answer to this question, leveraging our results on the calibration and power of SCEPTRE to guide our analysis. Consider a given perturbation-gene pair. Let *n*_trt_ be the number of cells containing the targeting perturbation. We refer to the number of cells containing *both* the targeting perturbation *and* nonzero expression of the gene as the “effective sample size” or the “number of nonzero treatment cells” for that pair. The SCEPTRE software filters out pairs for which the effective sample size falls below some threshold *T*. (By default, *T* is set to 7, a choice informed by the calibration (Figure S4 - S5) and power (Figure S8) results of SCEPTRE at different effective sample sizes.) Finally, let *s* denote the sparsity of a given dataset, i.e., the fraction of entries in the gene-by-cell expression matrix equal to zero.

We can model the effective sample size *X* of a given pair using a binomial model:

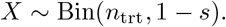

Among the *n*_trt_ cells that contain the perturbation, each exhibits nonzero expression of the gene with probability 1 *− s*. Thus, the number of cells containing *both* the perturbation *and* nonzero expression of the gene — i.e., the effective sample size — is binomially distributed with parameters *n*_trt_ and 1 *− s*. The probability that *X* equals *x* ∈ {0, …, *n*_trt_} is given by the binomial pmf:

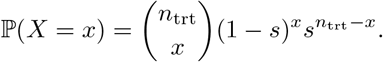

The pair passes pairwise QC only if *X* ≥ *T*, i.e., the effective sample size is greater than or equal to the pairwise QC threshold. Let *π* be the probability that the pair passes pairwise QC. (One should think of *π* as a parameter specified by the experimenter, like 0.9.) Finally, let *n*_trt_(*π, T, s*) be the number of treatment cells required for the pair to pass pairwise QC with probability *π* at pairwise QC threshold *T* on a dataset of sparsity *s*. We can express *n*_trt_(*π, T, s*) mathematically as

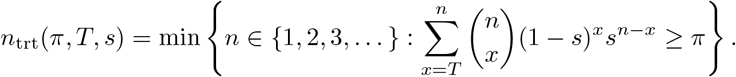

In other words *n*_trt_(*π, T, s*) is the smallest integer *n* such that the right-tail probability of Bin(*n*, 1*−s*) evaluated *T* is greater than or equal to *π*. It is challenging to derive an exact expression for *n*_trt_(*π, T, s*), but we can evaluate *n*_trt_(*π, T, s*) numerically. In summary, suppose that the sparsity level of the dataset is *s*, that the pairwise QC threshold is set to *T*, and that the desired probability that a given pair passes pairwise QC is *π*. Then the experimenter should ensure that each perturbation is contained within at least *n*_trt_(*π, T, s*) cells.

We evaluated *n*_trt_(*π, T, s*) for different values of *π, T*, and *s*, plotting *n*_trt_ as a function of *π* (Figure S13). We set the sparsity level *s* by computing the sparsity of the datasets analyzed in this work. The four “standard” datasets with gene expression readout — i.e., Frangieh co-culture, Frangieh control, Frangieh IFN-*γ*, and Papalexi (gene modality) — exhibited sparsity levels of 0.76, 0.77, 0.80, and 0.77, respectively (mean sparsity level 0.78). Next, the two TAP-seq datasets — i.e., Schraivogel chromosome 11 and Schraivogel chromosome 8 — exhibited sparsity levels of 0.33 and 0.27, respectively (mean sparsity level 0.30). Finally, the Papalexi (protein modality) dataset exhibited a sparsity level of 0.0. Thus, we set the sparsity *s* to 0.77, 0.30, and 0.0, corresponding to a “standard” screen with gene expression readout, a TAP-seq screen with gene expression readout, and a screen with protein expression readout.

We plotted the number of required treatment cells *n*_trt_(*π, T, s*) as a function of the probability *π* that a given pair passes pairwise QC (Figure S13). The left (resp., right) panel shows *n*_trt_(*π, T, s*) for an effective sample size QC threshold *T* of 7 (resp., 14). (The default threshold that the SCEPTRE software uses is 7.) Finally, the colors within each plot indicate the different sparsity levels: 0.78 (red), 0.30 (purple), and 0.0 (blue). The visual Figure S13 can be used to help guide experimental design. For example, suppose that an experimenter is designing a screen in which she plans to sequence gene transcripts using the standard single-cell sequencing protocol. (That is, the experimenter is not using the TAP-seq protocol.) Based on prior single-cell CRISPR screens datasets, the experimenter expects the sparsity of the dataset to be about 0.78. Moreover, suppose that the experimenter intends to use the default SCEPTRE pairwise QC threshold of *T* = 7. Finally, suppose that experimenter aims for 95% of the pairs to pass pairwise QC. Then the experimenter should try to ensure that each perturbation is contained within at least *n*_trt_(0.95, 7, 0.95) = 50 cells. As a very rough, general guideline, experimenters probably should ensure that each perturbation is contained within at least 50-80 cells. We (i.e., the authors) intend to add functionality to the SCEPTRE software in the future to aid users in designing single-cell CRISPR screen experiments, including a module for determining the approximate number of cells required per perturbation.

We note that the simple model considered here does not account for more complex phenomena present in single-cell CRISPR screen data. For example, cells containing multiple perturbations typically are removed as part of cellwise QC in low MOI, thereby decreasing the sample size. Moreover, sparsity can vary considerably across genes, as some genes are more highly expressed or more deeply sequenced than others. We believe that the analysis above constitutes a solid starting point for reasoning about experimental design in the context of SCEPTRE; however, additional further investigation remains to be done in a followup work.

## Footnotes

1 One potentially could make more discoveries using MIMOSCA by increasing the tuning parameter *B*, i.e. the number of times that the dataset is permuted and the test statistic recomputed. However, doing so would considerably increase computational cost.

2 The `wilcox.test` function on which Seurat-Wilcox relies returns *p*_exact_ only if (i) there are fewer than 50 cells across both treatment and control groups and (ii) no two cells (in either the treatment or the control group) have the same normalized expression level. This condition is expected to hold rarely, if ever.

3 A Cholesky decomposition of *Z*^*T*^ *WZ* could be used in place of the spectral decomposition, but the spectral decomposition is slightly more general, as it applies to matrices with eigenvalues equal to zero, which can occur (for example) when columns of *Z* are highly correlated.

1 There are several types of GLM residuals, including response (or “raw”) residuals, Pearson residuals, deviance residuals, and standardized (or “internally studentized”) residuals. We examined the simplest type of residual — namely, the raw residual — in this analysis. We internally performed comparisons to the other types of residuals as well, and we obtained broadly similar results (not shown).

2 We conjectured that the *T*_resid_-based permutation test controlled type-I error due to the CAMP phenomenon.

3 See timothy-barry.github.io/sceptre-book/run-calibration-check.html for details.

## Notes

https://www.dropbox.com/sh/jekmk1v4mr4kj3b/AAAhznGqk-TIZKhW40xiU6ORa?dl=0

### Competing Interest Statement

The authors have declared no competing interest.

### Summary of Updates

New supplementary materials section including several new analyses.

